# A fucose-binding superlectin from *Enterobacter cloacae* with high Lewis and ABO blood group antigen specificity

**DOI:** 10.1101/2024.10.01.616118

**Authors:** Ghamdan Beshr, Asfandyar Sikandar, Julia Gläser, Mario Fares, Roman Sommer, Stefanie Wagner, Jesko Köhnke, Alexander Titz

## Abstract

Bacteria frequently employ carbohydrate-binding proteins, so-called lectins, to colonize and persist in a host. Thus, bacterial lectins are attractive targets for the development of new antiinfectives. To find new potential targets for antiinfectives against pathogenic bacteria, we searched for homologs of *Pseudomonas aeruginosa* lectins and identified homologs of LecA in *Enterobacter* species. Here, we recombinantly produced and biophysically characterized a homolog that comprises one LecA domain and one additional novel protein domain. This protein was termed *Enterobacter cloacae* lectin A (EclA) and found to bind L-fucose. Glycan array analysis revealed a high specificity for the LewisA antigen and the type II H-antigen (blood group O) for EclA, while related antigens LewisX, Y, and B as well as blood group A or B were not bound. We developed a competitive binding assay to quantify blood group antigen binding specificity in solution. Finally, the crystal structure of EclA could be solved in complex with methyl α-L-selenofucoside. It revealed the unexpected binding of the carbohydrate ligand to the second domain, which comprises a novel fold that dimerizes via strand-swapping resulting in an intertwined beta sheet.

## Introduction

Bacterial infections are increasingly threatening as a consequence of antimicrobial resistance (AMR) development against standard-of-care antibiotics which have been on the market for many decades. Therefore, new treatment options are needed to fight the inevitable spread of multi-drug resistant pathogens rendering current medicines ineffective.(1) The World Health Organization has classified especially Gram-negative bacteria from the resistance-prone ESKAPE group as outstandingly important due to a lack of treatments with new modes of action, able to circumvent established antimicrobial resistance.(2)

For the critical priority Gram-negative bacterium *Pseudomonas aeruginosa*, many new approaches are actively being studied, such as new classes of antibiotics and alternative approaches interfering with bacterial virulence and pathogenicity.(1, 3) Among the so-called pathoblockers or anti-virulence agents(4, 5), interference with biofilm formation - a major determinant of drug resistance - is of particular interest to restore the pathogen’s susceptibility to treatments.(6) Numerous bacteria employ carbohydrate-binding proteins, so-called lectins, to adhere to host tissue and establish and maintain biofilms.(7–9) It has been demonstrated, that the *Pseudomonas* lectins LecA and LecB can be targeted with defined carbohydrates, small glycomimetics and dendrimers to prevent biofilms and interfere with bacterial virulence.(10–21)

Inspired by these data, we searched for orthologs of LecA in other bacteria. We previously identified many LecA orthologs in the genomes of several *Photorhabdus*, *Xenorhabdus* and *Enterobacter* species and experimentally characterized PllA from *Photorhabdus luminescens*, an insect pathogen.(22)

The *Enterobacter cloacae* complex is part of the ESKAPE pathogens infecting humans and comprises numerous species classified within 12 genetic clusters.(2, 23, 24) Bacteria associated with this complex are usually found in soil, sewage, drinking water reservoirs and are also part of the human intestinal microbiota.(25) *E. cloacae* and *E. hormaechei* constitute the most frequently isolated *Enterobacter* species from human clinical specimens.(24) Outbreaks of *E. cloacae* and the trigger to change from a commensal bacterium to a virulent one can be sporadic and often happens in the intensive care units of hospitals and causes either localized infections such as lung, wound, central nervous system, urinary tract, and catheter or orthopedic implant-associated infections, or systemic infections such as bacteraemia and sepsis.(25, 26) It is also well-established that *E. cloacae* is a culprit of outbreaks and fatalities in neonatal intensive care units of hospitals.(27) In their virulence arsenal, *E. cloacae* species employ many virulence factors like enterotoxins, α-hemolysin and thiol-activated pore-forming cytotoxins similar to Shiga-like toxin II after adhesion to epithelial cells.(28) In some clinical strains of *E. cloacae,* type III secretion system was also seen to be employed to destroy phagocytes and epithelial cells to facilitate host colonization.(29) Further, a type VI secretion system also showed to be instrumental in biofilm formation and adherence to epithelial cells.(30) Like other Gram-negative bacilli, the virulence of *E. cloacae* also depends on the presence of its outer membrane lipopolysaccharide which can help in avoiding opsonophagocytosis or initiate an inflammation cascade in the host cell leading to sepsis.(31) *E. cloacae* are multidrug resistant due to their chromosomally encoded and induced and/or constitutively-expressed AmpC β-lactamase with increased resistance rates following treatment with either β-lactams or 1st, 2nd and 3rd generation cephalosporins.(31) Fourth generation cephalosporins may be suitable against the AmpC β-lactamase strains if extended-spectrum β-lactamases (ESBL) are not present, in which case a combination therapy of colistin or aminoglycosides and carbapenems in a double regimen is used.(31) Due to the prevalence of ESBL and carbapenemases in this species, *E. cloacae* has become the third most common broad-spectrum Enterobacteriaceae involved in nosocomial infections, following *Escherichia coli* and *Klebsiella pneumoniae*.(32) For these reasons, the World Health Organization identified carbapenem-resistant Enterobacteriaceae (CRE) as a critical priority on its list of antibiotic-resistant bacteria in 2017, highlighting the urgent need for the development of new antibiotics. The characterization of LecA homologs in *Enterobacter spp* as possible anti-virulence targets is therefore of interest.

Here, we report the identification and biophysical and structural characterization of the first two-domain ortholog of LecA, found in *Enterobacter cloacae* subsp. *cloacae (*type strain: ATCC 13047), an important member of the *Enterobacter cloacae complex.* This strain was isolated from human cerebrospinal fluid and is the first completely sequenced member of the *E. cloacae* species. It possesses many virulence properties, encodes more than 50 antibiotic resistance genes, and has been extensively studied and used as a reference strain.(33–36)

The lectin termed EclA consists of an N-terminal LecA domain and a C-terminal domain reminiscent of carbohydrate binding modules. In this work, we established the L-fucose binding of EclA via its C-terminal domain while we did not succeed in identifying a ligand for the LecA-homologous N-terminus despite intense efforts. EclA forms homodimers resulting in the presentation of two N-termini towards one end and two C-termini towards the opposite end in its structure. This orientation suggests a function as a cross-linker for two carbohydrate ligands, one of which remains elusive. The unprecedented high specificity of EclA’s C-terminal domain for L-fucosides in mammalian blood group H-type II and LewisA antigens over related A/B-antigens or isomeric LewisX suggests a link to host binding specificity and pathophysiology.

## Results

Recently, we identified various orthologs of LecA in the genomes of diverse Gram-negative bacterial species including the insect pathogens *Photorhabdus luminescens* and *Xenorhabdus spp.*, and the opportunistic human pathogen *Enterobacter spp..*(*22*) Here, we have further analyzed the sequences of LecA orthologs from the ESKAPE pathogen *Enterobacter spp.*. Like LecA, most of them are short proteins comprised of only one LecA domain. Surprisingly, in the genome of the human spinal cord infection isolate of *Enterobacter cloacae subsp. cloacae* we identified a fusion protein, which we termed *Enterobacter cloacae* lectin A, EclA. Its gene *eclA* encodes for the protein EclA (ECL_04191, GenBank: ADF63724.1) composed of 283 amino acids with a molecular weight of 30.9 kDa. The protein sequence alignment of EclA with its orthologs LecA and PllA shows that the N-terminal domain of EclA is similar to the PA-1L (or LecA) family domain present in LecA and PllA (Figure 1). This domain spans from Trp20 to Glu133 and the amino acid involved only in calcium binding is conserved, while those involved in galactose-binding in LecA and PllA are not conserved. The second, C-terminal domain of EclA is recognized as a carbohydrate binding domain by bioinformatics sequence analysis tools.

**Figure 1:**
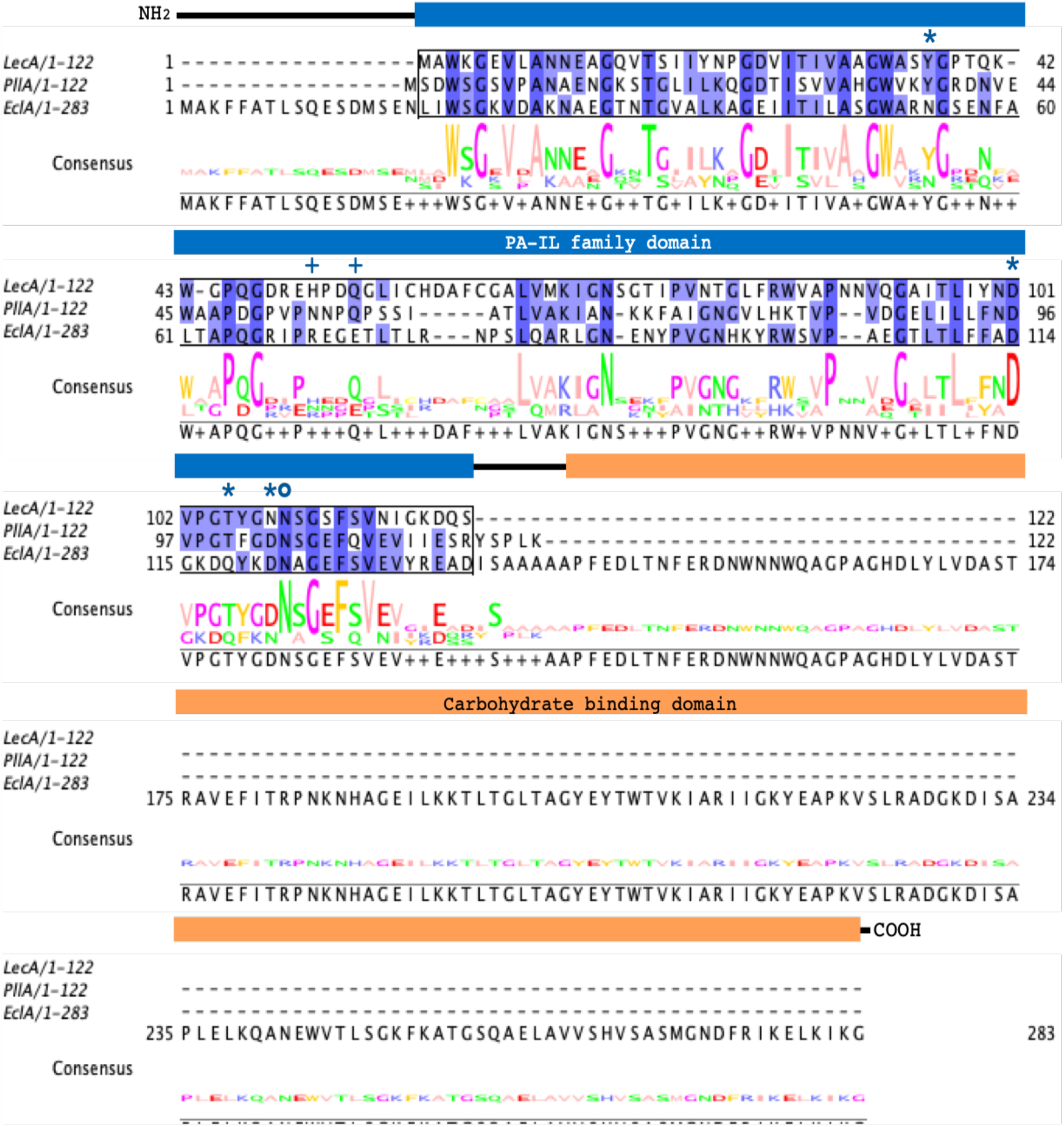
Sequence alignment of LecA, PllA and EclA. Protein sequences of LecA from *P. aeruginosa* PAO1, PllA from *P. luminescens subsp. laumondii* TTO1 and EclA from *E. cloacae subsp. cloacae* ATCC13047 were aligned in a multiple sequence alignment. The PA-1L family domain is indicated as blue box and the second domain of EclA is indicated as orange box. Amino acids are colored according to percentage identity (the higher the identity, the darker the color). Asterisks: Amino acids involved in ligand and calcium binding; open circle: amino acid involved in calcium binding; crosses: amino acids involved in ligand binding.

Since EclA is the first example of a LecA lectin domain fused to another domain, we set out to analyze this protein in depth. Full length EclA was cloned into the pET22 vector for recombinant cytosolic expression of the native protein in *E. coli*. The protein was highly expressed and present in the soluble fraction after cell lysis. The apparent molecular weight of EclA corresponded to the predicted value of 30.9 kDa as determined by denaturing gel electrophoresis (Figure S1). Despite the fact that LecA and PllA can both be purified on D-galactose-modified affinity resins, EclA did not bind to this resin and analogous purification failed. Therefore, EclA was finally purified from the soluble cell extract fraction by gel filtration on a superdex matrix (Figure S1).

### Carbohydrate binding specificity

To determine the ligand binding specificity of EclA, we analyzed its thermal denaturation in the presence of different carbohydrates (Figure 2). Among a set of 10 different monosaccharides tested, a detectable shift in EclA’s melting temperature (T_m_ = 55.2 °C) was only observed in presence of the deoxyhexose L-fucose (T_m_ = 57.2 °C), whereas all nine other carbohydrates including D-galactose did not influence EclA’s melting point. The result that EclA bound to L-fucose, i.e. 6-deoxy-L-galactose, was surprising as LecA and PllA bind to its hydroxylated enantiomer D-galactose. EclA was subsequently purified on a sephadex affinity resin modified with L-fucose and a yield of 30 mg EclA was obtained per liter bacterial culture (Figure S2).

**Figure 2:**
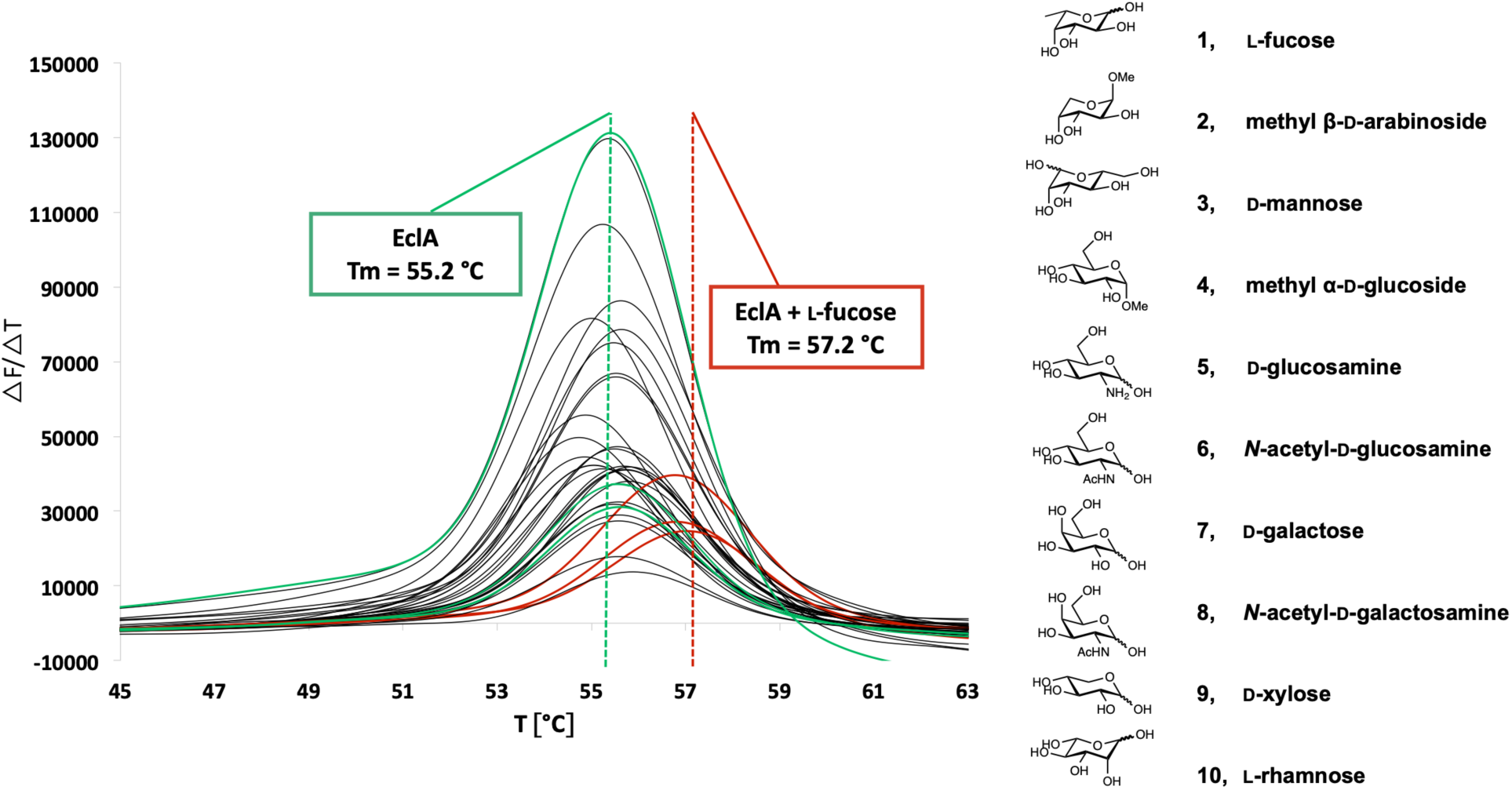
Thermal Shift Analysis of EclA ligand binding specificity indicates its binding to L-fucose.

To gain deeper insights into the carbohydrate binding specificity, we tested EclA on a glycan array. To this end, EclA was fluorescein-labelled with FITC and analyzed on the Consortium for Functional Glycomics’ Core H mammalian glycan array with 585 different carbohydrate epitopes (Figure 3). Binding of EclA was detected for a range of fucosylated oligosaccharides and the monosaccharide L-fucose on the array. In addition, the binding of EclA-FITC to sulfated sialyl-LacNAc was detected, the only bound ligand without a fucose residue.

**Figure 3:**
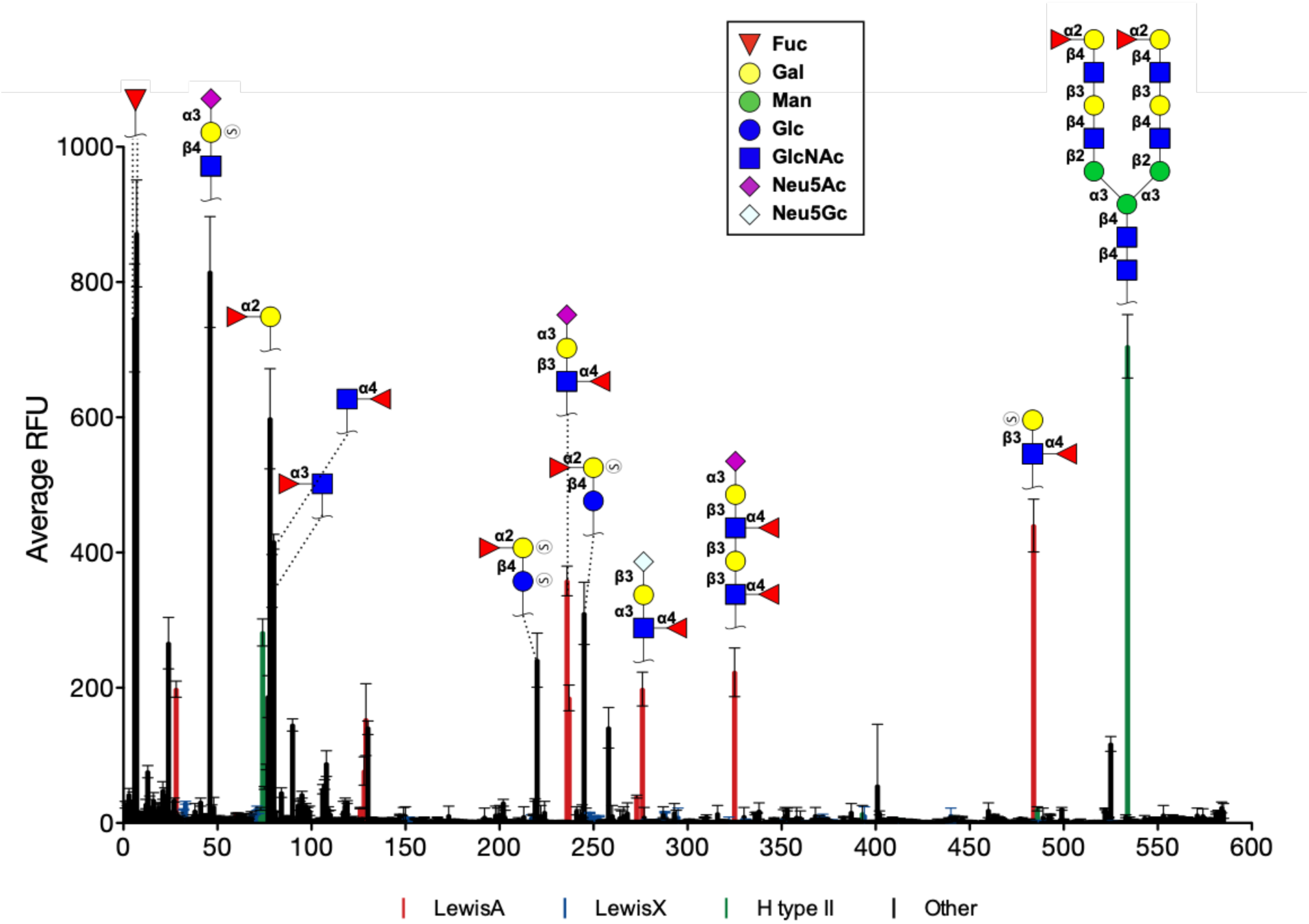
Glycan specificity analysis of fluorescein-labelled EclA on the CFG Core H glycan array. (S) = sulfate.

Careful analysis of the glycan array binding data for EclA revealed that fucosylated oligosaccharide ligands recognized by EclA were highly specific for only the blood group antigens LewisA and the H-antigen on a type II core (Figure 4). The selectivity of EclA for the LewisA antigen (Fuc-α-1,4-(Gal-β-1,3-)-GlcNAc) over its regioisomeric antigen LewisX (Fuc-α-1,3-(Gal-β-1,4-)-GlcNAc) is particularly remarkable. Additional sialylation was tolerated and LewisA and sialyl LewisA are generally recognized with high apparent affinity. Interestingly, out of 10 LewisX and 8 sialyl LewisX structures on the array, only ligand no. 24 carrying two additional sulfate residues was strongly recognized. Mono-sulfated epitopes were weakly recognized and no non-sulfated epitopes were bound. While α-2,3-sialylation of LewisA or LewisX has no influence on the specificity of EclA, an additional fucosylation resulting in LewisB and LewisY, respectively, abolished binding also for LewisB, a congener of LewisA.

**Figure 4:**
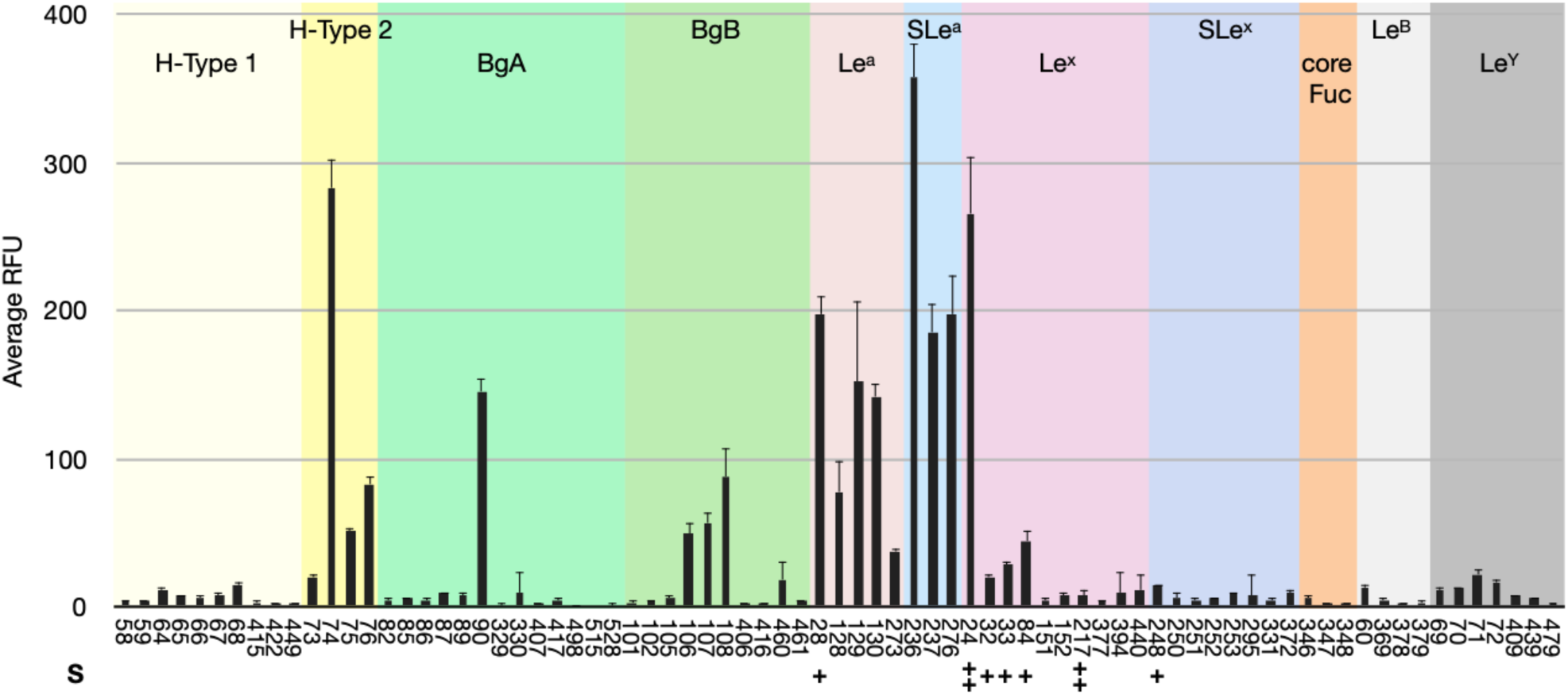
Analysis of blood group epitope specificity of EclA on the CFG Core H glycan array. Only unique and singly/monovalently displayed epitopes are compared here. Single (+) or double (++) sulfation of carbohydrate epitopes is indicated.

The observed selectivity of EclA for the H-type-II antigen (Fuc-α-1,2-Gal-β-1,4-GlcNAc) over isomeric H-type-I (Fuc-α-1,2-Gal-β-1,3-GlcNAc) is an interesting observation since the structural difference between those epitopes is located rather remote from the recognized fucose moiety, i.e., in the linking position of galactose to the GlcNAc residue. Blood group epitopes A or B are only occasionally recognized and a clear pattern is absent.

As mentioned, some carbohydrate ligands were recognized by EclA on the array that were sulfated. An analysis of all sulfated ligands on the array revealed the known specificity pattern and 3’-sulfated LewisA (no. 28, 484) and di(6,6’)- and mono(6 or 6’)- sulfated H-type II-like antigens (reducing end GlcNAc replaced by Glc, no. 220, 245, 258) were recognized by EclA. Among all sulfated glycans on the chip, only two more ligands were recognized: the above mentioned disulfated LewisX (no. 24) and the only binder devoid of fucose, 3’-sialyl-6’-sulfo-LacNAc (no. 46) carrying also two negative charges.

Initial crystallization attempts of full length EclA with a fucoside indicated ligand binding in the C-terminal domain (see below). Therefore, and to further dissect the binding specificities of the individual N- and C-terminal domains, we cloned and recombinantly produced them individually in *E. coli*: N-terminally His_6_-tagged EclA-N-terminus (termed EclA-N-tag, amino acids 2-140) and the native EclA C-terminus (termed EclA-C, amino acids 140-283).

Both protein constructs were subsequently purified to homogeneity using either Ni(II)-NTA affinity resin with imidazole elution or fucosylated-sepharose affinity resin with L-fucose elution (Figures S3, S4).

To quantify the binding of EclA with its carbohydrate ligands in solution, we developed a fluorescence polarization-based binding and displacement assay for EclA, in analogy to our previously described competitive binding assays for the bacterial lectins LecA(37), LecB(38, 39), PllA(22), BC2L-A(40) and BambL(41). Since EclA showed fucose-binding, the previously reported fluorescein-linked fucoside **11** was incubated with increasing concentrations of native EclA. Fluorescence polarization was monitored and fitting the obtained data resulted in K_D_s of 13.7 µM and 17.5 µM (two independent replicates) (Figure 5A). The experiment was also performed with EclA-C and a similar affinity was determined (K_D_ = 21.7 µM, Figure S5). Both constructs, EclA and EclA-C, were subsequently used to screen for inhibitors in a competitive binding assay (Figure 5B,C). The same monosaccharides previously used in the thermal shift assay were tested. In addition, methyl α- and β-L-fucosides were included to assess linkage specificity. The fucose-specificity from our previous assays was confirmed and only L-fucose (IC_50_ = 4.0-4.7 mM), methyl α-L-fucoside (IC_50_ = 1.3-1.4 mM) and its anomer methyl β-L-fucoside (IC_50_ = 1.5-1.8 mM) were inhibitors of, both, EclA and EclA-C. These data suggest that the hydrophobic methyl group of the aglycon is beneficial for binding, while α- or β-linkage shows no influence on binding.

**Figure 5:**
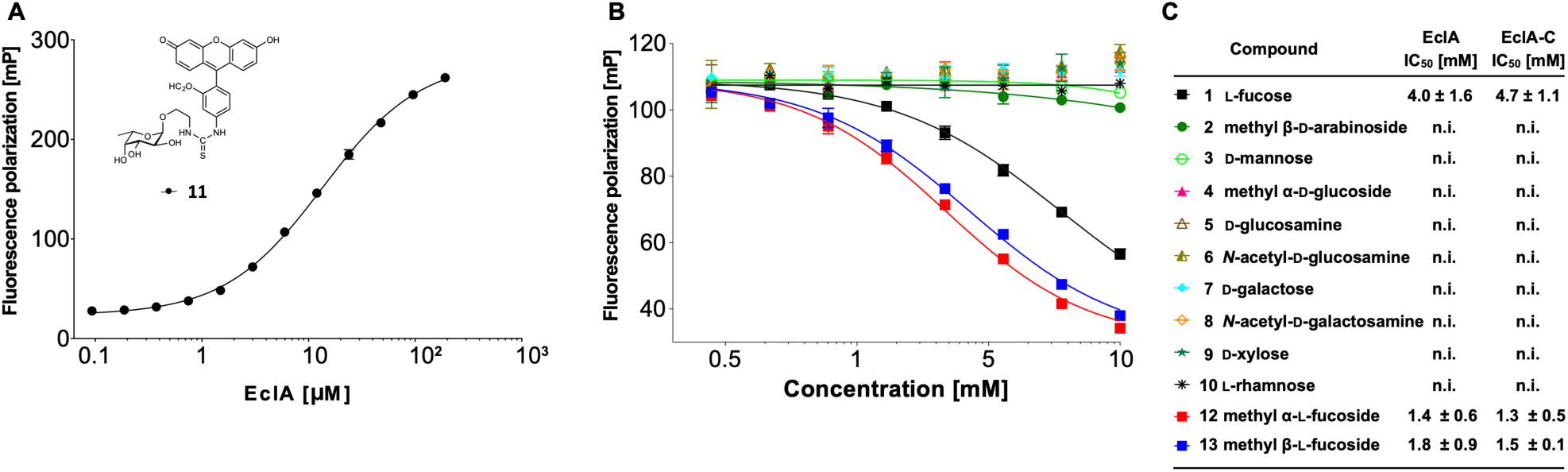
EclA ligand binding analyzed by fluorescence polarization. Direct binding of EclA to FITC-labelled fucoside **11** (A) and competitive inhibition (B) using a set of monosaccharides and methyl glycosides **1-10**, **12**, **13**. (C) IC_50_s and standard deviations for the inhibition of EclA or EclA-C derived from three independent titrations of triplicates each. Representative binding and inhibition data from one technical triplicate are depicted in (A) and (B).

Next, we also tested the human blood group antigens H-type I/II, A-type I/II, B-type-I/II, LewisA, LewisX, LewisB and LewisY in this competitive binding assay in solution (Figure 6). The data obtained corresponded to the semi-quantitative data obtained from the glycan array analysis: LewisA and H-type-II antigens were inhibitors of EclA and EclA-C with IC_50_s between 390 and 540 µM, whereas their isomeric epitopes LewisX and H-type-I showed an at least sevenfold weaker inhibition and the IC_50_s were at or above 3.75 mM.

**Figure 6:**
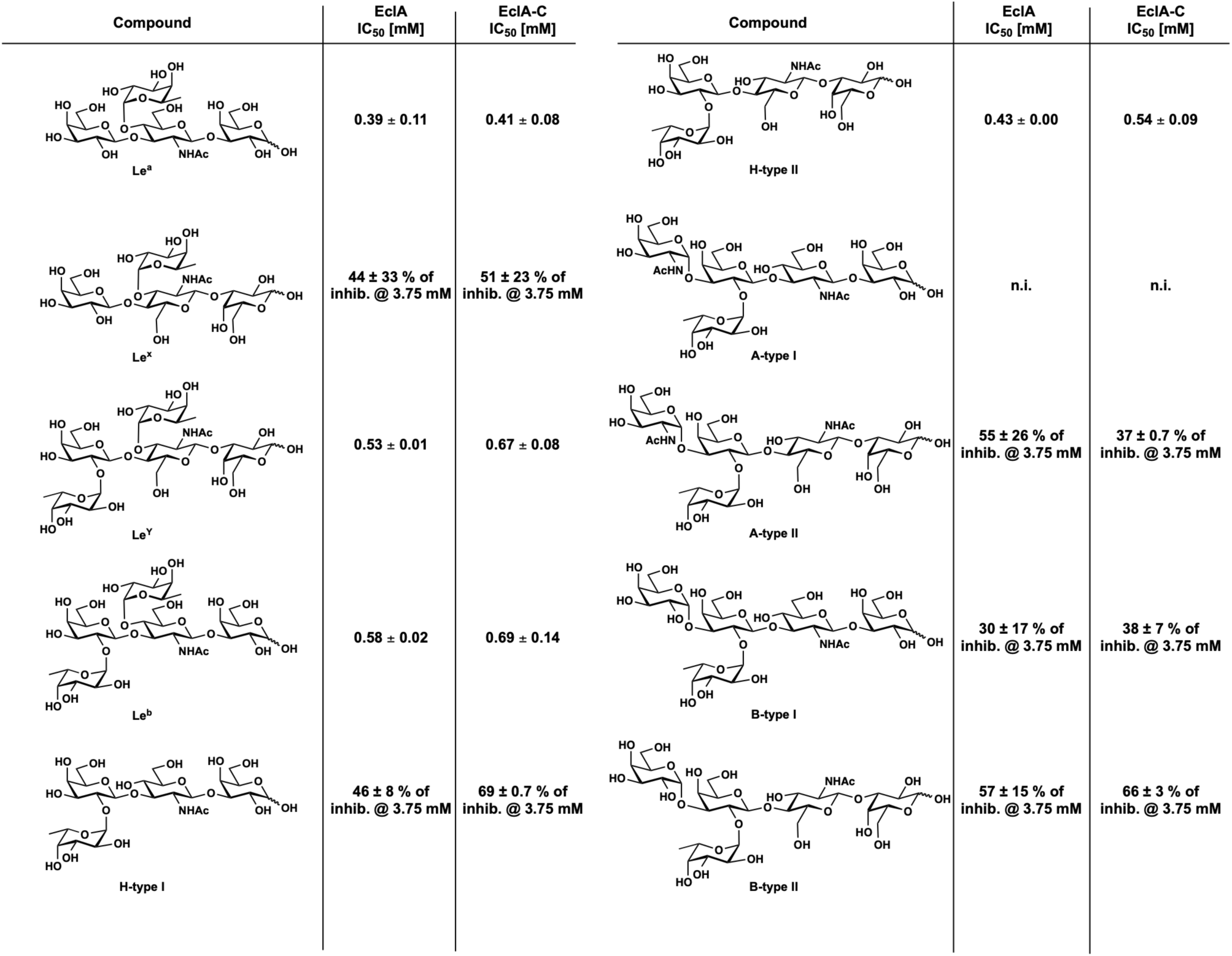
Competitive binding of human blood group antigens to EclA and EclA-C. IC_50_s and standard deviations are from three independent titrations of triplicates each.

In contrast to the surface binding in the glycan array, the difucosylated Lewis relatives LewisB and LewisY were also binders of EclA and its C-terminal domain in solution (IC_50_s between 530 and 690 µM). The blood group antigens A and B were only weak inhibitors of EclA and EclA-C with IC_50_s at or above 3.75 mM, which corresponds to the glycan array. Interestingly, we also observed an effect of type-II over type-I core.

To shed light on the ligand binding preference of the N-terminal domain of EclA, we tested full length EclA and both individual domains on a glycan array with a more diverse set of carbohydrate epitopes than the CFG mammalian glycan array. To this end, we labelled all three protein constructs with Cy3 and analyzed them on the Semiotik glycan array. This array contains 400 glycans and 200 bacterial polysaccharides. Full length EclA and its C-terminal domain showed binding specificity for fucose, LewisA and H-type antigens, while no ligand specific to the N-terminal domain of EclA could be identified, leaving its function enigmatic (Figure S6).

LecA and PllA are single domain proteins and form homotetrameric assemblies in solution which are also observed in their crystal structures.(22, 42) In contrast, their ortholog EclA consists of two separate domains, with one corresponding to a LecA domain. Thus, the elucidation of the oligomeric states of EclA is of interest. To this end, we analyzed EclA, EclA-N-tag and EclA-C by dynamic light scattering (DLS, Figure 7). The gene of EclA possesses two ATG codons at the 5’ end of the coding sequence which presumably led to the two protein species with very small apparent molecular weight differences as detected by SDS-PAGE (Figures S1, S2, S7). To test a homogeneous protein, we cloned the single amino acid variant of EclA-M14A, termed EclA-L, and expressed and purified it as reported for EclA (Figure S7). The obtained estimated molecular weights of the protein species in solution showed an observed molecular weight of 80.1 kDa for full length EclA-L (MW = 30.9 kDa), 43.5 kDa for the N-terminal domain (EclA-N-tag, MW = 16.0 kDa), and 33.0 kDa for the C-terminal domain (EclA-C, MW = 15.9 kDa). The observed molecular weight of EclA-C has a rather narrow distribution and suggests dimer formation. EclA-L and EclA-N-tag both have a larger dispersion of molecular weights with a value close to 2.5-fold of their calculated molecular weight. DLS measures the hydrodynamic radius and the molecular mass of the proteins is calculated assuming an ideal spherical particle, which explains these deviations. Taken together, all three constructs showed experimental values for their molecular weight between 2.1- and 2.7-fold of their calculated molecular weights, suggesting dimer formation in all three cases.

**Figure 7:**
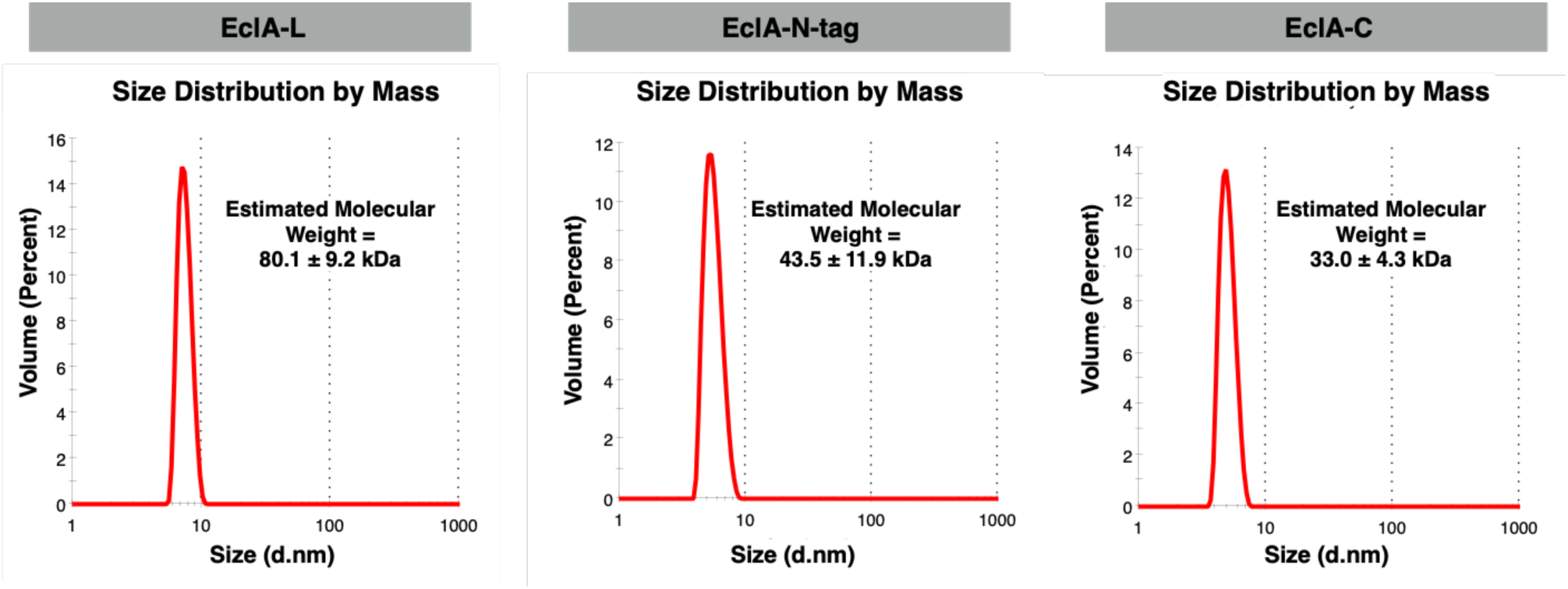
Dynamic light scattering analysis of EclA-L (EclA-M14A), EclA-N-tag and EclA-C.

### Crystallography

In order to shed light on structure and ligand binding of the two EclA carbohydrate recognition domains crystallographic studies were first carried out on full length EclA. We obtained crystals in several conditions that were all of similar morphology and diffracted well to beyond 2.5 Å. The datasets could be processed in P2_1_ with excellent statistics and showed no obvious abnormalities. Molecular replacement with Phaser(43) using a monomer of the lectin PllA (PDB ID 5ofz), which is homologous to the N-terminal half of EclA (EclA-N), resulted in a solution containing a dimer of the N-terminal domain, with the expected dimeric arrangement of lectin domains and a TFZ score of 12.3, indicating a correct solution. Unexpectedly, we were unable to resolve the C-terminal domain of EclA, even though the crystal packing of the N-terminal domain clearly left sufficient space for it. Extensive trials to refine the full structure failed and we thus resorted to treating the N- and C-terminal domains of EclA separately.

The crystal structure of EclA-N-tag was determined in space group P2_1_ and revealed dimers, which matched the partial solution obtained for the full-length protein (Figure 8). The overall structure of EclA-N-tag is highly similar to the structure of the *Photorhabdus luminescens* lectin PllA (Cα rmsd of 0.47 Å over 93 atoms; Figure 8B), with each monomer adopting a jelly-roll type β-sandwich fold as previously described.(22) Unambiguous electron density was observed for the canonical calcium ion at the putative carbohydrate-binding site, coordinated by the side chain oxygens of Asp114, Asp121, Asn122, and the main chain oxygens of Asn54 and Gln118 (Figure S8A). One structural difference that stands out between the two structures is the size of the ligand-binding pockets: it is much wider in EclA-N than in PllA, which is the result of a movement of a loop (residues 70 – 82; Figure 8B, 8C). This could be a crystallization artefact, as H-bond interactions between this loop and symmetry mates can be observed. An overlay of all four EclA-N molecules shows an almost identical orientation of this loop, which may indicate a biological significance (Figure S8B). This is supported by the involvement of residues present on the corresponding loops of other lectins in ligand-binding (Figure S8C). However, in the absence of any known carbohydrate-ligand it is difficult to ascertain the role of the observed wide pocket and the loop in ligand-binding specificity of EclA-N.

**Figure 8:**
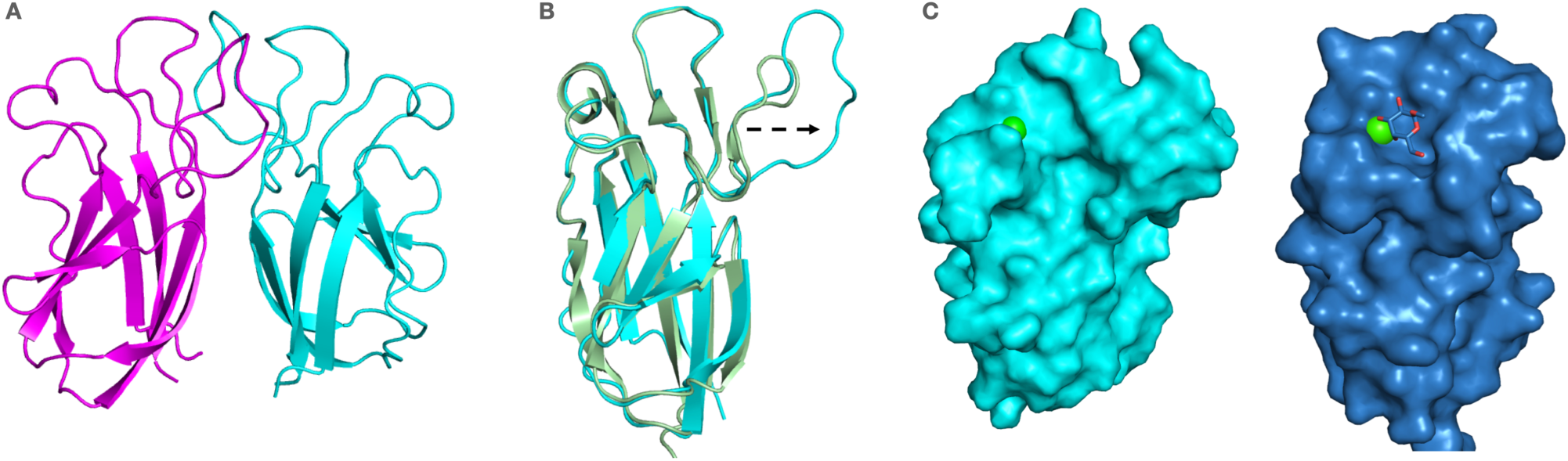
Crystal structure of the N-terminal domain of EclA (EclA-N-tag) and comparison with its ortholog PllA. A. Dimer of EclA-N-tag, B. superposition of the monomers of EclA-N-tag and PllA indicate the extended loop in EclA leading to opening of the putative binding site. C. Surface illustration of the crystal structure of the N-terminal domain of EclA (EclA-N-tag, left) and comparison with its ortholog PllA with bound galactose (right). Bound ligand is shown as stick and Ca^+2^ ions as green spheres.

The C-terminal construct, EclA-C, yielded very well diffracting crystals belonging to space-group P2_1_2_1_2_1_. Molecular replacement was unsuccessful; therefore, we expressed and crystallized selenomethionine labelled EclA-C. We were able to obtain an initial solution with a dimer in the asymmetric unit. Mirroring the issues of the full-length EclA construct, unambiguous, clear electron density could only be observed for one of the two dimers. We hypothesized that the intrinsic flexibility of the protein was the culprit. After collecting many datasets with different additives and ligands, we were able to obtain a dataset of EclA-C in complex with methyl α-L-selenofucoside (MeSeFuc) and were able to refine this structure fully. In the structure, this ligand appears to limit the flexibility of the dimer by cross-linking symmetry-mates via the carbohydrate ligand (MeSeFuc interacts with Glus178, Ile214, Ile215, and Arg275, Figure S9).

MeSeFuc is present in a small cleft, stabilized by a number of hydrophobic, metal and H-bond interactions (Figure 9). The cis-diol of the fucoside coordinates the calcium ion, which itself is tightly coordinated by Ile180, Thr181, Ser269, Gly271 and Asp273. In addition to the calcium coordination, the 4-OH of the ligand is in hydrogen bonding distance of the backbone NH of Gly271 and the side chain carboxylate of Asp273. The carbohydrate residue forms one additional hydrogen bond via its 2-OH and a bound water molecule with the side chain of Ser269. Furthermore, extended hydrophobic contacts of the aglycon and the glycosidic selenium are formed with Met270 and Tyr218. The exocyclic methyl group of the fucoside is accommodated by a hydrophobic pocket formed by Ile215, Lys217 and Tyr218.

**Figure 9:**
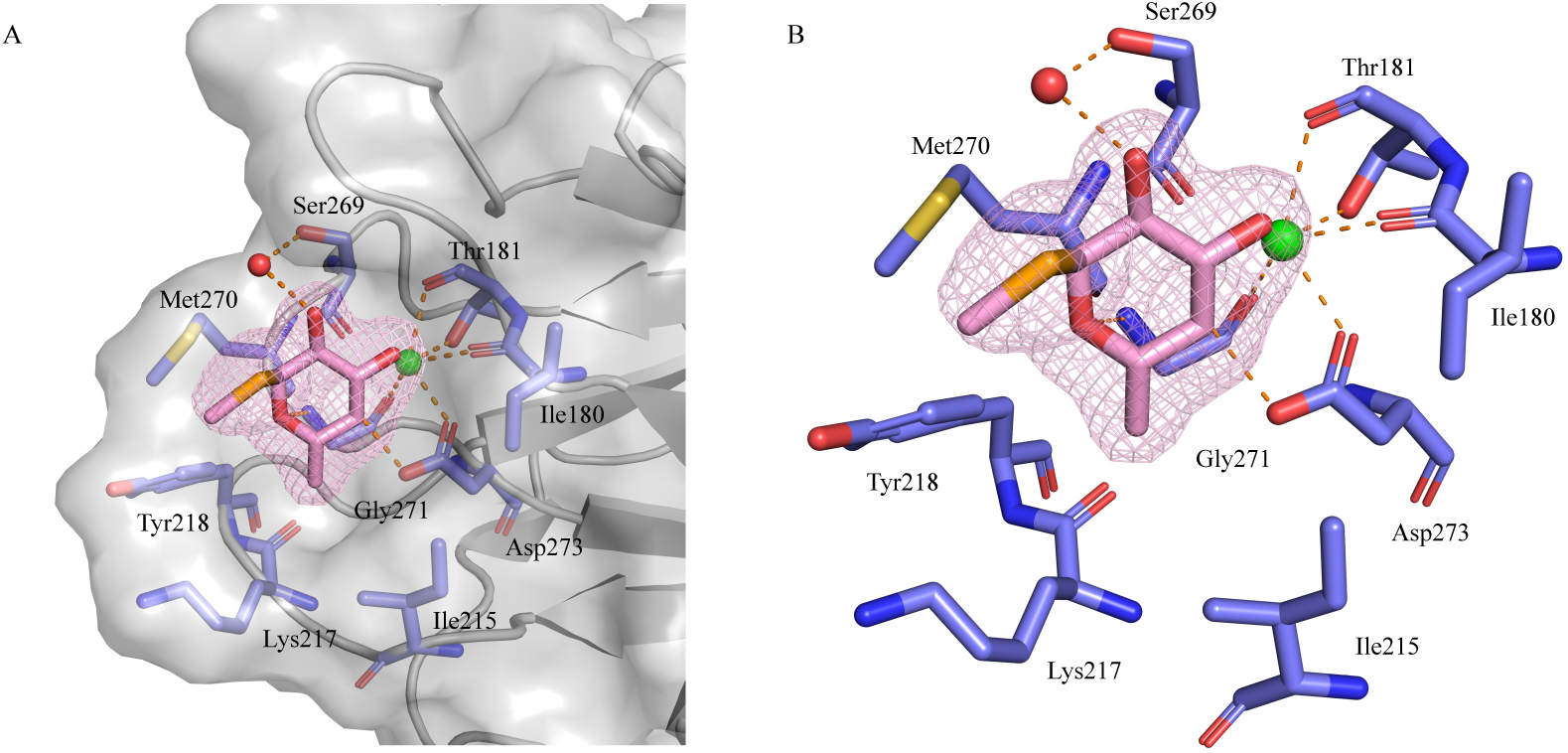
Crystal structure of EclA-C in complex with methyl α-L-selenofucoside. A. Surface display of the protein illustrating the ligand’s interactions in the binding pocket, B. Coordination of the protein-bound calcium ion and hydrogen bonding interactions of the fucoside. Electron density displayed as mesh at 4 σ.

The dimer partners of EclA’s C-terminus interact mainly via their N-termini through a β-strand swap, which results in a twisted anti-parallel β-zipper (Figure 10A and B). This mode of interaction presumably results in a very unusual arrangement in the full-length protein (Figure 11A). A search for structural homologs of EclA-C using the DALI server(44) revealed the carbohydrate binding module (CBM) CBM22-1 of *Clostridium thermocellum* endo-1,4-β-D-xylanase (Xyn10B, PDB ID 2W5F) to be the closest structural homolog (Z-score of 9.8; Figure 10C). Akin to CBM22-1, each monomer of the EclA-C structure consists of a β-sandwich fold, comprised of two anti-parallel β-sheets made from 7 β-strands (β4, β5, β6, β7, β9, β10, β11; CBM22-1 numbering). In EclA-C, β4 is extended and connected to an additional β-strand (β3) that is not part of the β-sandwich (Figure 10B and C; Figure S10).

**Figure 10:**
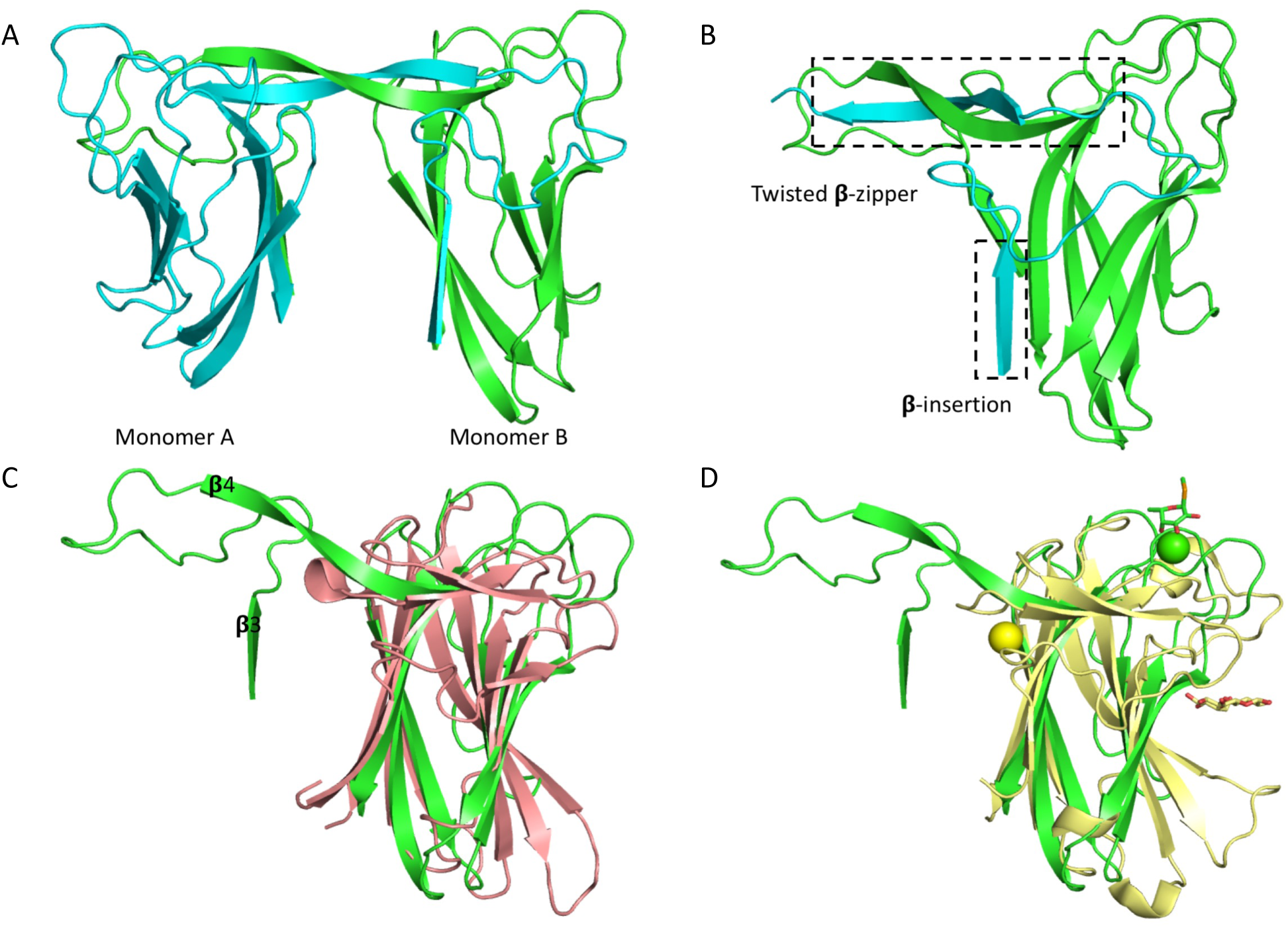
Structural analysis of the C-terminal domain of EclA (EclA-C). A. Dimer comprising the asymmetric unit is shown as ribbon model with the monomers coloured cyan and green. B. The twisted zipper formation and the additional β-insertion observed in the crystal structure of EclA-C. Only β-sheet (3 and 4) of monomer A are shown. C. Superposition of EclA-C structure (green) with CBM22-1 (PDB ID: 2W5F; salmon). Cα RMSDs approx. 3.6 Å over the entire length of the protein (monomer B). D. Superposition of EclA-C structure (green) with CBM22-2 (PDB ID: 4XUQ (yellow). Ligands are shown as sticks (methyl selenofucoside, green; and xylotriose, yellow) and the Ca^+2^ ions as spheres (EclA-C, green; and CBM22-2, yellow).

**Figure 11:**
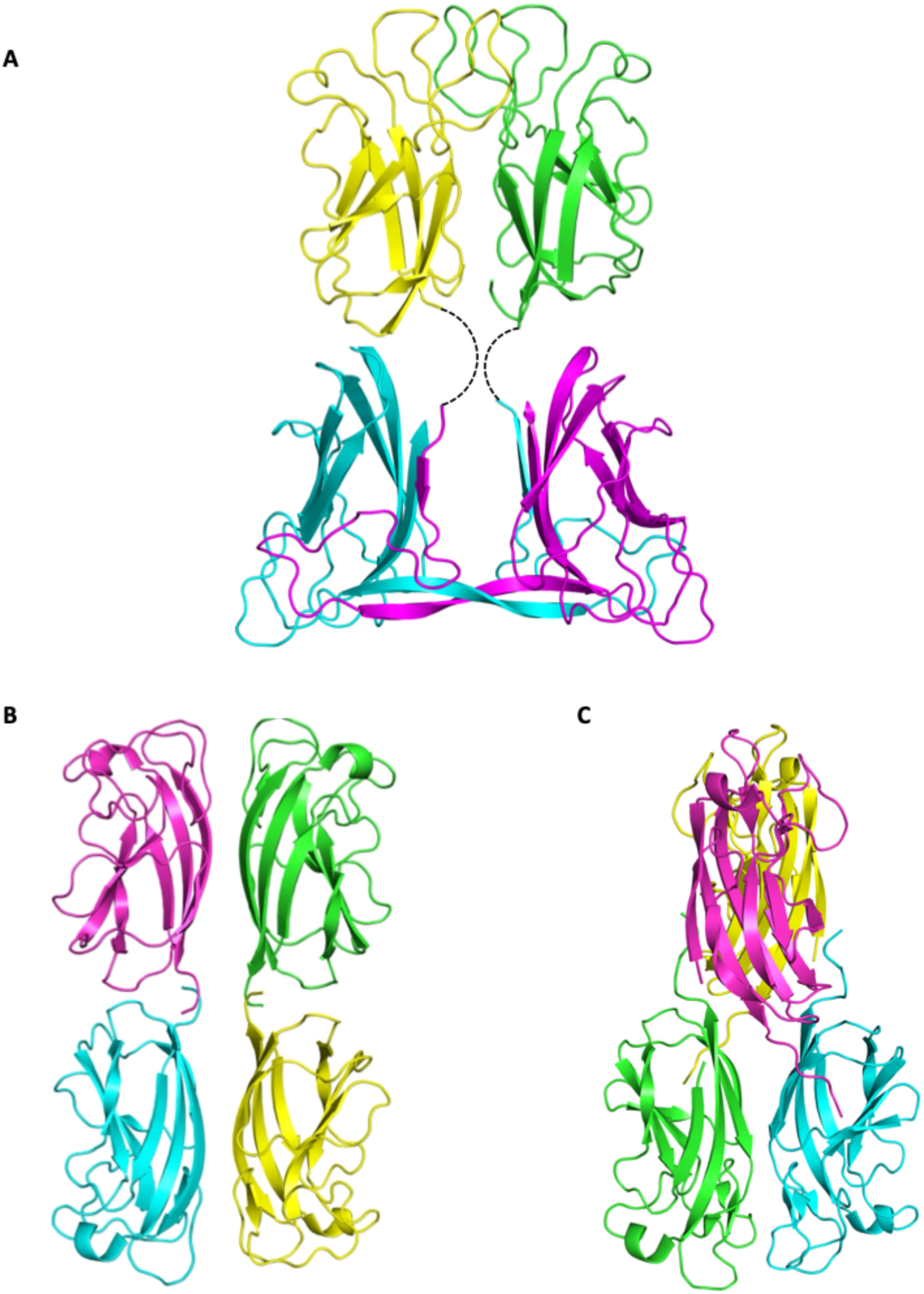
Comparison of quaternary structures of homodimer EclA with homotetramers LecA and PllA. A. Model of the quaternary structure of EclA using the separate crystal structures of EclA-C and EclA-N-tag. Dashed lines indicate the missing 11 amino acids between N- and C-terminus. B. Tetramer of LecA in its rectangular orientation (pdb code 4LKD(42)). C. Tetramer of PllA with both dimers twisted by 90° to each other (pdb code 5OFZ(22)).

Despite the overall similar structure, the observed mode of Ca^2+^ ion and ligand-binding in the structure of EclA-C is different compared other CBMs; the binding of the carbohydrate ligands is located at different areas of the protein and in the EclA structure primarily stabilized by the Ca^2+^ ion (Figures 9, 10D, and S11) rather than a network of H-bonds as observed for e.g., CBM67, CBM60 and CBM36. As a result, the removal of Ca^2+^ had a drastic effect of the EclA-C ligand binding. Moreover, CBMs often contain a second, structural Ca^2+^ ion, which is absent in the EclA-C structure.

### Bioinformatics

Following the discovery of EclA’s unique structure and ligand binding specificity, we next analyzed the abundance of EclA in other *Enterobacter* strains. The protein sequence of EclA from *E. cloacae* subsp*. cloacae* ATCC13047 was used as template in a tblastn search in *Enterobacter* genomes. Nearly 50 homologs of EclA have been identified and are shown in a multiple sequence alignment (Figure 12).

**Figure 12:**
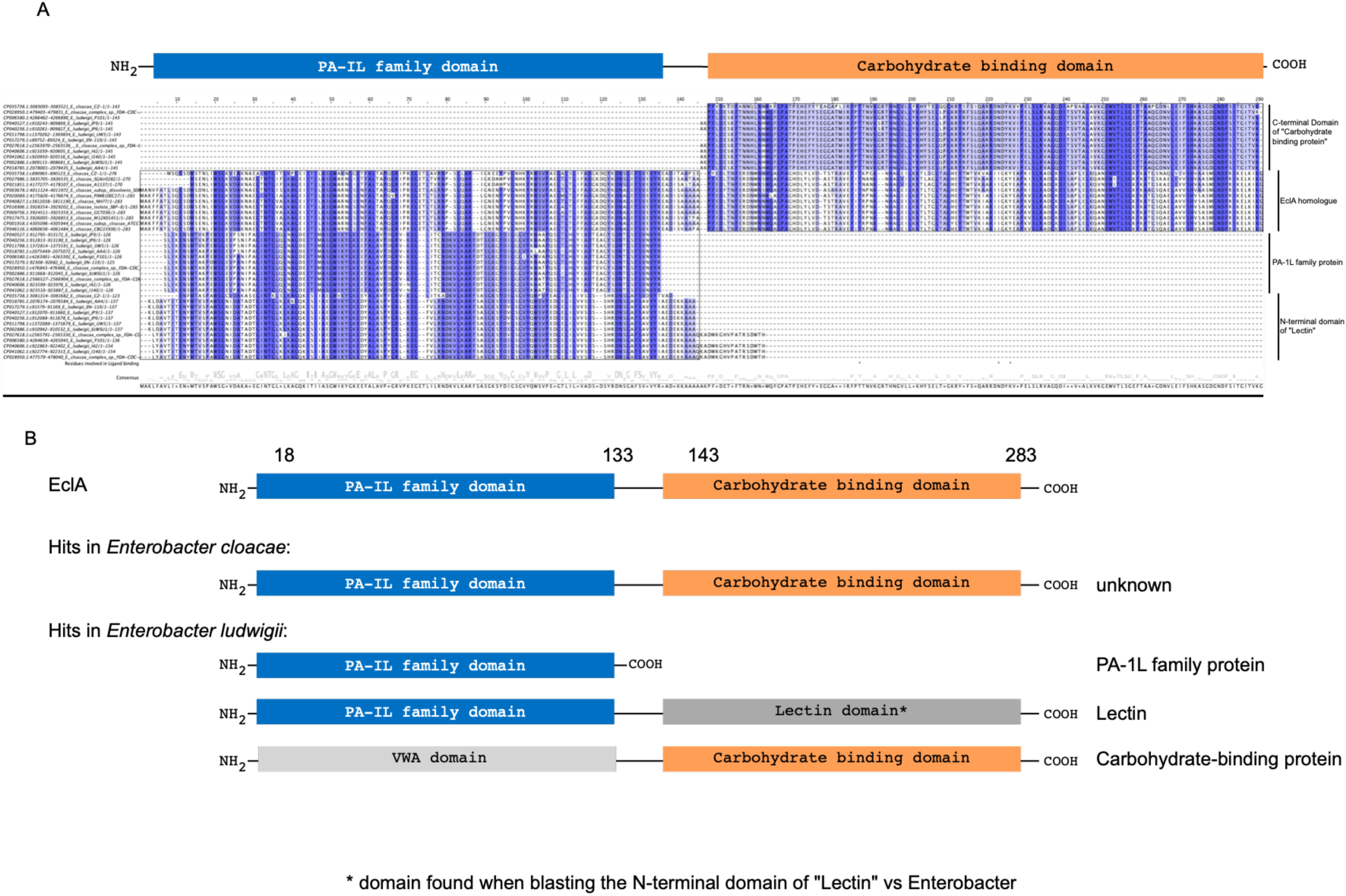
Multiple sequence alignment of EclA homologs in the *Enterobacter* taxid. A: Protein sequences of hits from a tblastn search were aligned in a multiple sequence alignment. PA-1L family domain is indicated as blue box and the C-terminal domain of EclA is indicated as orange box. Amino acids are labelled according to percentage identity (the higher the identity, the darker the color). For illustration purposes, found hits were truncated and only the sequence parts matching EclA were included in the alignment. B: Schematic representation of proteins found with the BLAST search. In *E. cloacae* genomes, only genes of unknown function were found. In *E. ludwigii* genomes, three proteins were identified that match either the N-terminal or the C-terminal domain of EclA: an annotated lectin, an annotated carbohydrate-binding protein and a member of the PA-1L protein family.

Interestingly, three different groups of homologs could be identified: (i) in *E. cloacae* strains, proteins of unknown function matching full-length EclA were identified (sequence identities: 99 - 85%); and in *E. ludwigii* strains, proteins harboring either one (ii) or the other (iii) of the two EclA domains were identified but sequences covering both domains simultaneously were absent (Figure 12B).

Moreover, each of the three complete *E. ludwigii* genomes harbors three proteins that match to one EclA domain (Figure 12B): (i) one single domain protein of unknown function annotated as a PA-1L family protein with 45-39% sequence identities to EclA-N; (ii) a protein of unknown function, annotated as lectin harboring two domains, of which the N-terminal domain shares 42-38% sequence identity with EclA-N; and (iii) another protein of unknown function annotated as carbohydrate-binding protein harboring two domains, of which the C-terminal domain shares 35-33% sequence identity with EclA-C. The C-terminal domain of the ‘lectin’ hits found in *E. ludwigii* genomes is similar to its N-terminal domain, while the N-terminal domain of the ‘carbohydrate binding protein’ is a von Willebrand Factor type A domain (vWA), named after its presence in von Willebrand Factor (vWF) - a large glycoprotein found in blood plasma. Interestingly, the genes found in *E. ludwigii* genomes encoding the three proteins that share either the LecA domain or the C-terminal domain of EclA are in direct proximity to each other on the genome.

In a second tblastn search on orthologs of the EclA C-terminal domain, the taxid *Enterobacteriaceae* was excluded. The results showed mainly *Pseudomonas* genomes (ca. 83%, but not *P. aeruginosa*) and some *Serratia* genomes (ca. 17%) carrying one gene, which is of unknown, function but annotated as !carbohydrate protein” (e.g., *Pseudomonas asplenii* strain ATCC 23835, LT629777.1, data not shown). The C-terminal domain of this protein shares a sequence identity with the C-terminal domain of EclA of 38-26%.

## Discussion

*Enterobacter* spp. belong to the critical ESKAPE pathogens. In this work we identified a new lectin, termed EclA, from *Enterobacter cloacae* as an ortholog of LecA from *P. aeruginosa*. Interestingly, EclA consists of two domains, an N-terminal LecA/PA-IL domain and a C-terminal carbohydrate binding domain. The N-terminal domain is also homologous to PllA from *P. luminescens* and both proteins, LecA and PllA, bind D-galactosides as ligands. To analyze EclA structure and ligand binding, we recombinantly expressed the protein and a first purification attempt on a galactose-affinity matrix surprisingly failed. Subsequent thermal shift analysis of EclA purified by gel filtration unexpectedly indicated the thermal stabilization of EclA in presence of L-fucose.

The binding specificity of EclA was analyzed on the mammalian glycan array of the Consortium for Functional Glycomics. Among the approximately 600 mammalian glycans, a highly specific binding preference of EclA could be established. In general, fucose-containing ligands were recognized by EclA, with an unusually high specificity for LewisA and H-type-II blood group antigens. Their isomeric structures LewisX and H-type-I were not bound. In addition, larger oligosaccharide analogs of LewisA and H-type II carrying additional fucose or Gal/GalNAc residues, such as LewisB and blood group A and B antigens were also not bound. Interestingly, one additional oligosaccharide was bound by EclA that did not have a fucose residue but is negatively charged due to sulfation and sialylation.

We then cloned and expressed the genes of the individual domains of EclA to analyze their binding properties independently. The fucose-binding was unequivocally assigned to the C-terminal domain. Solution phase binding preferences have been established by fluorescence polarization using a fluorescence-labelled fucoside and various ABO and the Lewis human blood group antigens. The competitive binding assay confirmed the high specificity of EclA for LewisA over isomeric LewisX and for the H-type II antigen over isomeric H-type I antigen oligosaccharides.

In an attempt to identify the ligand of the N-terminal domain, EclA and its two individual domains were then further analyzed on a complementary glycan array from Semiotik comprising approximately 400 glycans and 200 polysaccharides. In general, this array also reproduced the established ligand binding preferences determined for the full length EclA. The Semiotik array further confirmed EclAߣs preference for fucose, LewisA and H-type II antigens located at the C-terminal domain, while unfortunately no specific ligand could be identified for the N-terminal domain, the ortholog of the galactose-binding relatives LecA and PllA.

To uncover the protein’s structure, we then crystallized EclA first as the full-length protein and then both individual domains. The structure of the full-length protein crystals could unfortunately not be solved for both domains, in contrast to the structures of the individual domains. A small fucoside ligand was coordinated via its 3,4-dihydroxy cis-diol motif via one calcium-ion by the C-terminal domain. Additional hydrogen-bonds and lipophilic contacts of the ligand with EclA explain the binding specificity for fucose-containing ligands. Interestingly, the C-terminal domain formed dimers in the crystal with a very unusual dimerization mode via their N-termini through a β-strand insertion and a twisted anti-parallel β-zipper.

The LecA domain located at the N-terminus of EclA also formed dimers in the crystal structure with very high overall similarity to PllA/EclA. The major difference concerns a loop located in the binding site of LecA and PllA that participates in galactoside binding in both proteins. This loop is extended in EclA, leading to a large opening of the site. In addition, three arginines form an extended positively charged patch at the surface adjacent to the conserved calcium-ion that is mediating galactose-binding in LecA and PllA. This positively charged area might explain the binding to several charged carbohydrates that were not recognized by the C-terminal domain as their neutral congeners (e.g., sulfated LewisX or sulfated sialyl LacNAc), although not all charged carbohydrates were recognized and a clear understanding of these observed binding is still missing. Further analysis for the identification of the binding specificity of the EclA N-terminal domain remains a challenge.

Lectins are often multivalent to increase binding strength through avidity. The spatial presentation of binding sites in lectins is important for their function. For surface binding, lectins present their carbohydrate-binding sites on the same side as in Shiga toxin which invades gut epithelial cells after binding to glycolipids on a cell(45), whereas the interaction with two or more partners involves the presentation of their binding sites in opposing directions which allows cross-linking of glycosylated binding partners as observed by the native hemagglutination of red blood cells for many lectins and demonstrated by AFM for the crosslinking of artifical glyco-systems by LecA(46).

Our data from X-ray crystallography and DLS strongly suggest that the structure of EclA orients the total four carbohydrate binding sites in opposite directions, in analogy to its homologs, LecA and PllA. However, since LecA and PllA are homotetramers assembled as dimers of dimers of the same short single domain polypeptide, they present four identical binding sites. In contrast, EclA is a dimer of a two-domain polypeptide chain resulting in the presentation of two identical domains to either side. This orientation suggests the cross linking of two distinct binding partners by EclA, one of which has been elucidated in this work, i.e., the blood group antigens LewisA and H-type II.

In summary, we have presented a novel lectin termed EclA from the opportunistic human pathogen *Enterobacter cloacae*. The two-domain protein forms homodimers with an unusual structural assembly. While its C-terminal domain is specific for LewisA and H-type II oligosaccharides, ligand binding specificity of its LecA-homologous N-terminal domain remains enigmatic. The ortholog LecA is a known virulence factor for infections with *P. aeruginosa* and has been shown to bind to host glycolipids for cell entry. LecA is furthermore a key factor for biofilm formation. Thus, LecA has become a drug target for fighting infections with the high priority pathogen *P. aeruginosa*.(12, 19) Whether EclA has an analogous role in infection of the ESKAPE member *E. cloacae* remains to be elucidated. Still, its mode of action will be more complex due to the two distinct domains.

*Enterobacter* species are frequently isolated from neonatal sepsis.(24) Blood groups have been widely analyzed and these cell surface carbohydrates are primary receptors for infections by viruses and bacteria.(47, 48) Interestingly, Lewis A is overrepresented in neonates. It is known that newborns are LewisA positive and during childhood most individuals develop into LewisB positive, a ligand that is recognized with much weaker affinity by EclA. LewisA is strongly expressed in the gastrointestinal tract, the urinary tract and in the lungs: these are the prime loci for infection with the *E. cloacae*. In contrast epitopes of the ABO blood group system are expressed broadly across the body. Interestingly, Boral et al. reported that “the group O neonates had significantly more sepsis due to *Enterobacter cloacae*”.(49) Therefore, EclA could potentially be a link between *Enterobacter* infections and neonatal sepsis, which merits further analysis. Furthermore, the type I core is present on tissue oligosaccharides while type II core is found in secreted soluble blood group antigens.(50) Possibly, EclA evaded its inhibition by soluble oligosaccharides and evolved specificity for the tissue resident type I core structure for efficient infection.

## Materials and Methods

Methyl ⍺-L-fucoside (**12**) and methyl β-L-fucoside (**13**) were purchased from Carbosynth (Oxford, UK). Methyl ⍺-D-glucoside (**4**), *N-*acetylgalactosamine (**8**), *N*-acetylglucosamine (**6**), D-xylose (**9**) were obtained from Sigma Aldrich Chemie (Germany); L-fucose (**1**), D-mannose (**3**), and D-galactose (**7**) from Dextra Laboratories (Reading, UK); D-glucosamine (**5**) from Calbiochem (USA); methyl β-D-arabinoside (**2**) from TCI Deutschland GmbH (Germany); L-rhamnose (**10**) from AppliChem (Germany); Histo-blood group antigens were purchased from Elicityl OligoTech (France).

Fluorescent ligand FITC-fucoside **11** was synthesized as described.(38) Methyl α-L-selenofucoside was synthesized according to the previously published protocols from fucose.(51, 52)

### Cloning, expression, and purification of recombinant EclA, EclA-L, EclA-N-tag, EclA-C

The synthetic gene of EclA (ECL_04191, GenBank: ADF63724.1) was cloned tag-free into the vector pET22b and the resulting plasmid was purchased from Eurofins Genomics (Germany). The gene sequences of EclA-N-tag and EclA-C were amplified by PCR using Phusion polymerase (New England Biolabs, UK) with the designed primers containing restriction sites and N-terminal His6-tag sequence if needed as listed in Table S1. The protein sequence of EclA is MAKFFATLSQESDMSENLIWSGKVDAKNAEGTNTGVALKAGEIITILASGWARNGSENFALTAPQG RIPREGETLTLRNPSLQARLGNENYPVGNHKYRWSVPAEGTLTLFFADGKDQYKDNAGEFSVEVYREADISAAAAAPFEDLTNFERDNWNNWQAGPAGHDLYLVDASTRAVEFITRPNKNHAGEILKKTLTGLTAGYEYTWTVKIARIIGKYEAPKVSLRADGKDISAPLELKQANEWVTLSGKFKATGSQAELAVVSHVSASMGNDFRIKELKIKG (283 aa, MW 30.9 kDa, calculated pI = 6.54). The protein sequence of EclA-L, where a second possible start codon at position 14 is mutated (M14A), in order to have only one isoform of EclA is MAKFFATLSQESDASENLIWSGKVDAKNAEGTNTGVALKAGEIITILASGWARNGSENFALTAPQGRIPREGETLTLRNPSLQARLGNENYPVGNHKYRWSVPAEGTLTLFFADGKDQYKDNAGEFSVEVYREADISAAAAAPFEDLTNFERDNWNNWQAGPAGHDLYLVDASTRAVEFITRPNKNHAGEILKKTLTGLTAGYEYTWTVKIARIIGKYEAPKVSLRADGKDISAPLELKQANEWVTLSGKFKATGSQAELAVVSHVSASMGNDFRIKELKIKG (283 aa, MW 30.9 kDa, calculated pI = 6.54). The protein sequence of EclA-N-tag, based on EclA-L (see below) is MHHHHHHAKFFATLSQESDASENLIWSGKVDAKNAEGTNTGVALKAGEIITILASGWARNGSENFALTAPQGRIPREGETLTLRNPSLQARLGNENYPVGNHKYRWSVPAEGTLTLFFADGKDQYKDNAGEFSVEVYREADISAAA (146 aa, MW 16.0 kDa, calculated pI = 5.95), the protein sequence of EclA-C is MAAPFEDLTNFERDNWNNWQAGPAGHDLYLVDASTRAVEFITRPNKNHAGEILKKTLTGLTAGYEYTWTVKIARIIGKYEAPKVSLRADGKDISAPLELKQANEWVTLSGKFKATGSQAELAVVSHVSASMGNDFRIKELKIKG (144 aa, MW 15.9 kDa, calculated pI = 8.8).

After digestion of the expression vector pET22b (+) (Novagen, Germany) and the PCR product with NdeI and BamHI (New England Biolabs, UK), ligation of the insert was performed using TakaRa DNA ligation kit (TAKARA Bio Inc, Japan) resulting in plasmid pET22b*-eclA-N-tag* and pET22b-*eclA-C*. The sequences were confirmed by sequencing (GATC Biotech, Germany) with primers T7 promotor and T7 terminator (Table S1).

For expression, pET22b-*eclA*, pET22b-*eclA-N-tag*, and pET22b-*eclA-C* were transformed into chemically competent *E. coli* BL21(DE3), and the expression strain was selected on LB agar supplemented with ampicillin (100 µg mL^-1^). 2 liters of LB supplemented with ampicillin (100 µg mL^-1^) were inoculated with a preculture and grown at 37 °C and 180 rpm to OD_600_ of 0.5 – 0.6. Expression was induced by addition of IPTG (0.15 mM final concentration), and bacteria were then further cultured for 4 h at 30 °C and 180 rpm. The cells were harvested by centrifugation (9000 g, 10 min), and the pellet was washed with TBS/Ca buffer (20 mM Tris, 137 mM NaCl, 2.6 mM KCl, pH 7.4, supplemented with 100 µM CaCl_2_). The cells were resuspended in 40 mL of TBS/Ca supplemented with PMSF (1 mM) and lysozyme (0.4 mg mL^-1^) and subsequently disrupted by five cycles on a SONOPULS ultrasonic homogeniser (BANDELIN electronic, Germany). Cell debris was removed by centrifugation (19000 g, 60 min), and the supernatant was loaded onto a gel filtration column (see below) or a column containing fucose-coupled sepharose CL-6B (for EclA and EclA-C). The affinity resin was prepared using L-fucose in analogy to the procedure reported by Fornstedt and Porath.(53) The column was washed with TBS/Ca buffer. EclA and EclA-C were then eluted by addition of 100 mM L-fucose to the TBS/Ca buffer. The eluted fractions were extensively dialyzed against distilled water and then TBS/Ca buffer. For the His-tagged protein (EclA-N-tag), the supernatant was loaded on a 5 mL HiTrap affinity column (GE Healthcare, Germany). The column was washed with TBS/Ca supplemented with 20 mM imidazole and then EclA-N-tag was eluted by addition of 250 mM imidazole to the TBS/Ca buffer. The eluted fractions were thereafter desalted on HiTrap desalting column (GE Healthcare, Germany). The concentration of EclA, EclA-N-tag, EclA-C were determined by UV absorbance at 280 nm using calculated molar extinction coefficients of 50420, 22460, 27960 M^-1^ cm ^-1^, respectively. The yield of purified EclA, EclA-N-tag, EclA-C was 15, 5, 12 mg per liter of culture volume, respectively.

EclA-L (M14A) was generated by QuikChange site-directed mutagenesis(54) using plasmid pET22b-*eclA* as the template and phusion polymerase (New England Biolabs, UK). The PCR forward and reverse primers GB32 and GB33 (Table S1, MWG-Biotech AG, Ebersberg, Germany) were designed to introduce point mutation at the desired position. The reaction for either a forward or a reverse primer was separately performed in a 25 μL volume using a PCR thermocycler (Professional thermocycler, Analytik Jena, Germany). Thereafter, the separately amplified DNA strands were combined and reannealed to give pET22b*-eclA-L*. The parent template was digested by DpnI restriction enzymes (New England Biolabs, UK) (> 1 h at 37 °C) and then 10 μL of reaction was transformed (electroporation, 1800 V, 25 μF, 200 Ω, 5.2 ms) into *E. coli* XL1-blue cells and plated onto LB agar supplemented with 100 µg mL^-1^ ampicillin. After overnight incubation at 37 °C, a clone was selected, grown in liquid culture and the plasmid isolated (GenEluteTM Plasmid miniprep Kit, SIGMA, Germany). Finally, the isolated plasmid was confirmed by sequencing at GATC (Konstanz, Germany).

### Gel filtration

A HiLoad 16/600 Superdex 200 pg column (GE Healthcare) was equilibrated with TBS/Ca buffer (20 mM Tris, 137 mM NaCl, 2.6 mM KCl, pH 7.4, supplemented with 1 mM CaCl_2_) at a flow rate of 1 mL/min. A calibration curve was used from our previous study. Thereafter, EclA was loaded on the column and analyzed with the same flow rate.

### Dynamic light scattering (DLS) measurements

DLS measurements were performed on a Zetasizer Nano-ZS (Malvern Instruments, UK). Stock solutions were filtered with a syringe filter before measurements. Either 100 µM of EclA-L or 200 µM of EclA-N-tag and EclA-C in TBS/Ca (20 mM Tris, 137 mM NaCl, 2.6 mM KCl, pH 7.4, supplemented with 1 mM CaCl_2_) was measured at 25 °C.

### Fluorescence labeling of EclA constructs and glycan array analysis

Protein (1.4 mL of 60 µM EclA in Na_2_CO_3_ buffer, pH 9.3) was incubated at room temperature under shaking (500 rpm) with fluorescein isothiocyanate (FITC, 66 µL of 3 mg mL^-1^, in sodium carbonate buffer, pH 9.3) for 1 h. Purification of the labeled protein was performed as described above for unlabeled EclA; the protein concentration was determined as described previously for LecB-PA14(39) using an extinction coefficient 50420 M^-1^ cm^-1^.

FITC-labeled EclA was tested on the Consortium for Functional Glycomics (CFG) mammalian glycan array (Core H) version 5.3 containing 585 printed glycans in replicates of 6. Standard procedures of Core H (details see https://research.bidmc.org/ncfg) were run at 5 and 50 µg mL^-1^ protein based on the protocol by Blixt *et al.*(*55*). Raw data of the EclA binding experiments at 5 and 50 µg mL^-1^ are available as in Table S3.

For analysis on the Semiotik glycan array, EclA-L, EclA-C, and EclA-N-tag were individually labelled using Cy3-NHS ester (Toronto Research Chemicals, Canada). Briefly, 700 µg of protein in sodium carbonate buffer (190 µL, pH 8.3) were mixed with 10 µL Cy3-NHS ester solution (4.7 mg/mL in DMSO). The reaction was incubated for 4 h at r.t. under exclusion of light. The labelled proteins were desalted and excess dye was removed in spin columns using TBS/Ca. The labelled proteins were used without further purification. The samples were analyzed on the Semiotik glycan array as described.(56) Array slides were incubated with protein samples at 50 µg/mL (in PBS supplemented with 0.05% Tween 20) for 1 hour at 37 °C and shaking (32-34 rpm). Slides were analyzed after washing on a PerkinElmer ScanArray Gx (gain 60, laser power 80, resolution 20 µm).

### Thermal Shift Assay

The thermal shift assay was conducted using a StepOnePlus Real Time PCR device (Applied Biosystems. USA). 10 µL of a solution of each tested compound at 20 mM in TBS/Ca was transferred in triplicate into white semi skirted RT-PCR 96-well plate (Axon-lab, Germany). Thereafter, 10 µL of a solution of 10 µM of EclA and 10X of SYPRO Orange dye in TBS/Ca was added to each well. The plate was centrifuged for 1 min at 4000 rpm and covered by RT-PCR 96-well plate foil (Axon-lab, Germany). The final concentration of EclA, SYPRO orange, and compounds were 5-10 µM, 5-10X, and 10 mM, respectively. The plate was heated from 20 to 99 °C with a heating rate of 0.5 °C min^-1^. Fluorescence was monitored at Ex/Em: 490/530 nm.

### Direct titration of fluorescent ligand **11** with EclA or EclA-C

A serial dilution of EclA or EclA-C (10 µL, 0.4 – 400 µM) was transferred in triplicates to a black 384-well plate (Greiner Bio-One, Germany, catalog no. 781900) and then 10 µL of 20 nM of fluorescent ligand **11** in TBS/Ca buffer was added to each well. The plate was centrifuged for 1 min at 4000 rpm. After incubation at r.t. for 1 hour, blank corrected fluorescence intensity was recorded using a PheraStar FS microplate reader (BMG Labtech GmbH, Germany) with excitation filters at 485 nm and emission filters at 535 nm, and fluorescence polarization was calculated. The data were analyzed using a four-parameter fit of the MARS Data Analysis Software (BMG Labtech GmbH, Germany). A minimum of three independent experiments on three plates was performed.

### Competitive binding assay: Displacement of fluorescent ligand **11** from EclA, EclA-L or EclA-C

Each tested monosaccharide and the Histo-blood group antigens were serially diluted from 0.07 - 20 mM and 0.03 - 3.5 mM, respectively. Thereafter, 10 µL of each concentration was transferred to a 384-well plate (Greiner Bio-One, Germany, catalog no. 781900) in triplicates. Then, 10 µL of 20 µM of EclA or EclA-C and 10 nM of fluorescent ligand **11** in TBS/Ca was added to each well. The plate was centrifuged for 1 min at 4000 rpm. After incubation at r.t. for 1 hour, blank corrected fluorescence intensity was recorded using a PheraStar FS microplate reader (BMG Labtech GmbH, Germany) with excitation filters at 485 nm and emission filters at 535 nm, and fluorescence polarization was calculated. The data were analyzed using a four-parameter fit of the MARS Data Analysis Software (BMG Labtech GmbH, Germany). A minimum of two independent experiments on two plates was performed for each compound.

### Crystallization

Crystals of EclA-N (12 mg/mL) were obtained in 0.06 M divalents (magnesium chloride hexahydrate and calcium chloride dihydrate), 0.1 M Tris and bicine, pH 8.5, 40% v/v glycerol and 20% w/v PEG 4000. EclA-C (8 mg/mL) was preincubated with methyl α-L-selenofucoside (1-2 mM) prior to setting up crystallization trials. The optimized crystals were obtained in 100-280 mM disodium phosphate, 16-22% w/v PEG 3350, and cryoprotected with 30% v/v glycerol. Data were collected at SLS (EclA N-terminus; beamline X06DA) and ESRF (EclA-C terminus; beamline MASSIF-3). EclA-N-tag structure was solved using PHASER Molecular Replacement(43) with PllA (PDB ID 5OFZ) as a search model, whereas the EclA-C structure was solved using the anomalous signals from the Se-containing sugar. Data were processed using XDS(57) or Xia2(58) and the initial models obtained using PHENIX AutoSol(59) were rebuilt manually with COOT(60) and refined using PHENIX(61). Both structures were validated using MolProbity(62), and the images presented were created using PyMOL (Schrödinger, LLC) and Ligplot+(63). Protein crystallography data collection and refinement statistics are depicted in Table S2.

### Bioinformatics

The protein sequence of EclA from *Enterobacter cloacae subsp. cloacae* ATCC13047 was used in a tblastn search in the Enterobacter taxid using the Basic Local Alignment Search Tool (BLAST) on the NCBI/NLM/NIH website (https://blast.ncbi.nlm.nih.gov/Blast.cgi). In a second tblastn search, the taxid Enterobacter was excluded from the search and only the C-terminal domain of EclA was used as a query.

Sequence alignments were performed using the multiple sequence alignment tool Clustal Omega.(64) Amino acids are labelled according to percentage identity (the higher the identity, the darker the color) using Jalview(65) software.

## Acknowledgements

We acknowledge Dr. Eike Siebs (HIPS, Saarbrücken) for performing thermal shift assays on EclA-N-tag. The authors acknowledge excellent technical support from Dirk Hauck (HIPS, Saarbrücken). We further acknowledge the participation of the Protein-Glycan Interaction Resource of the CFG and the National Center for Functional Glycomics (NCFG) at Beth Israel Deaconess Medical Center, Harvard Medical School (supporting grant R24 GM137763). Dr. Jamie Heimburg-Molinaro, Dr. Richard D. Cummings from the Consortium for Functional Glycomics and Dr. Nadya Shilova from Semiotik LLC are gratefully acknowledged for performing and analyzing glycan arrays. A.T. acknowledges funding from the TANDEM graduate school of Saarland University. Support from EU COST Action EureStop is kindly acknowledged.

