## Supporting Information for "A fucose-binding superlectin from *Enterobacter cloacae* with high Lewis and ABO blood group antigen specificity"

4 current address: Leibniz University of Hannover

*corresponding authors

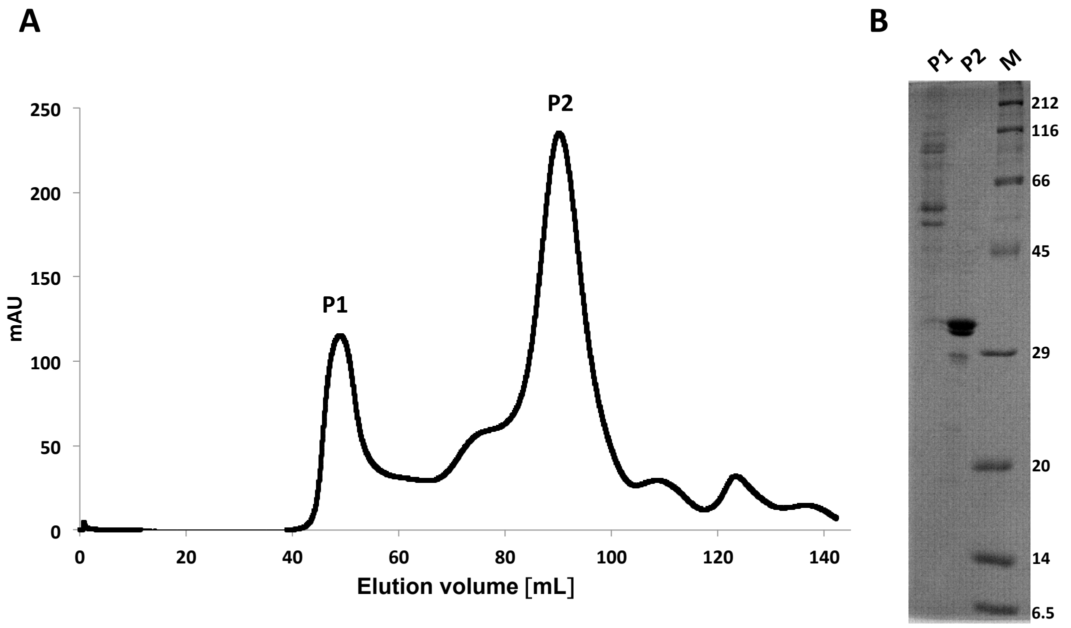


**Figure S1:** A. Gel filtration of culture supernatant producing EclA. B. SDS-PAGE of fraction P1 and P2 shows EclA present in P2.

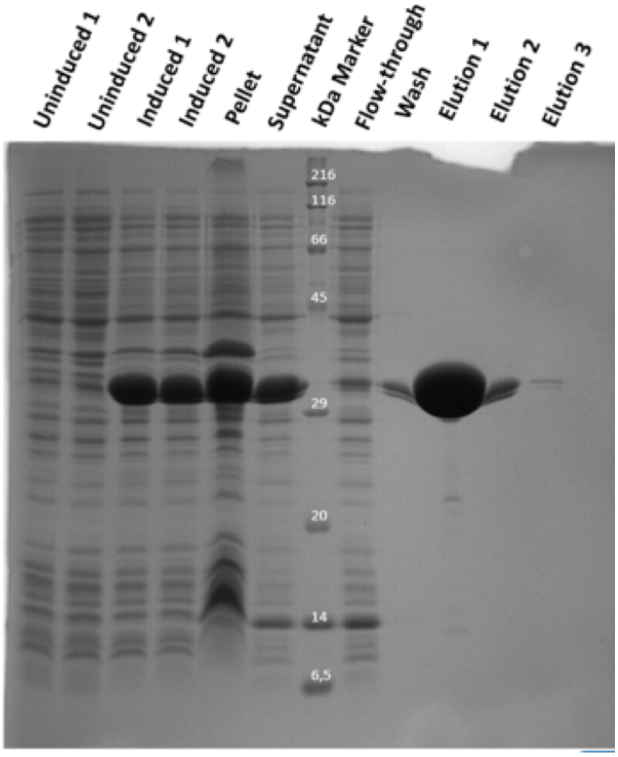


**Figure S2**: SDS-PAGE analysis of expression and purification of EclA on fucose-coupled sepharose resin with elution using 100 mM L-Fuc; yield: ca. 60 mg / 2 L

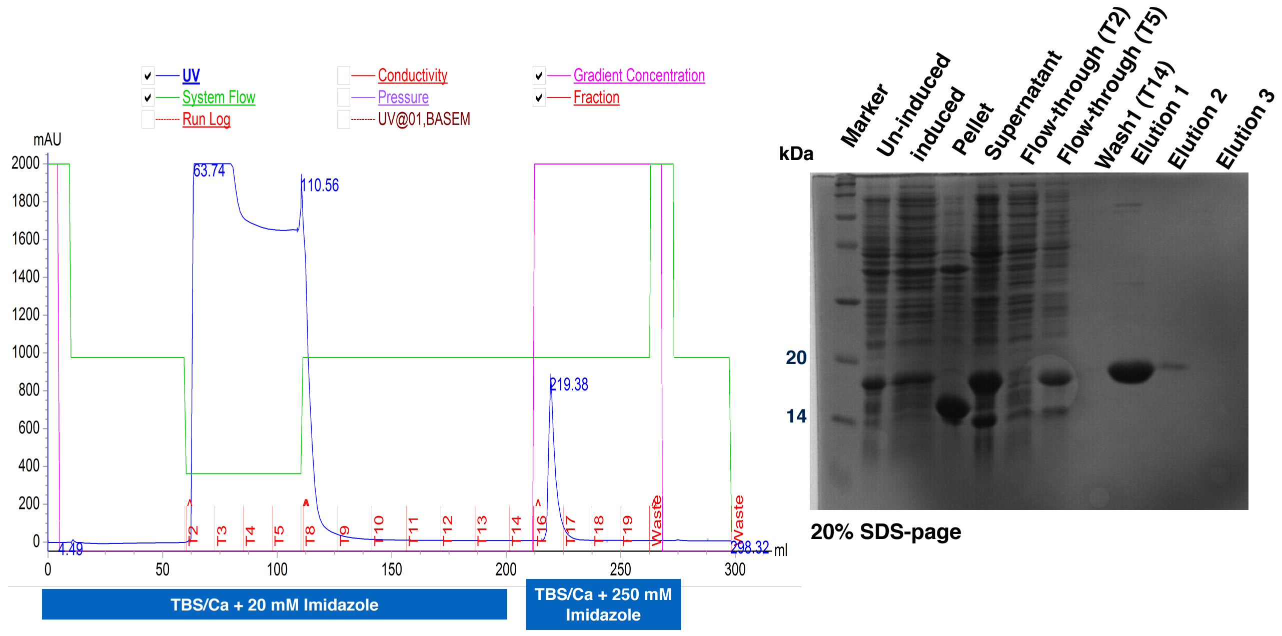


**Figure S3.** Expression and purification of EclA-N-tag on a HisTrap (Ni(II) sepharose) column. Left: the chromatogram of purification steps; right: 20% SDS-PAGE analysis of the expression and purification steps.

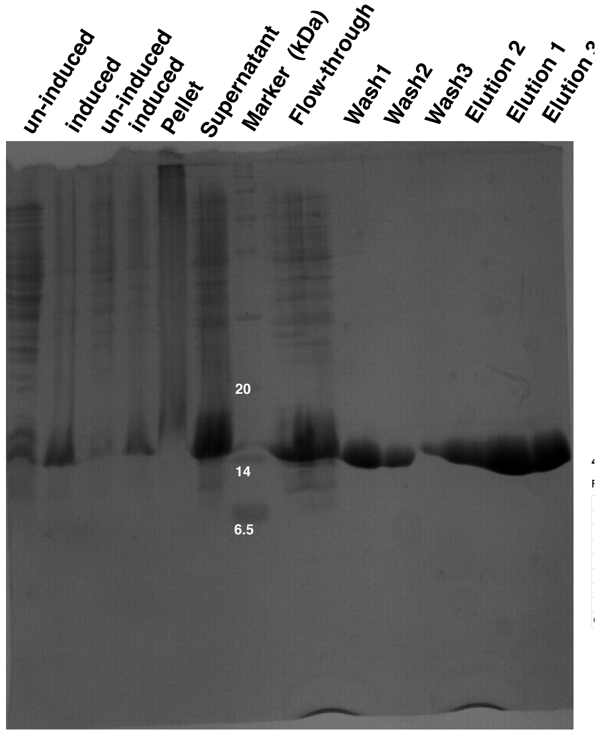


**Figure S4**. Expression and purification of EclA-C analyzed by 20% of SDS-PAGE.

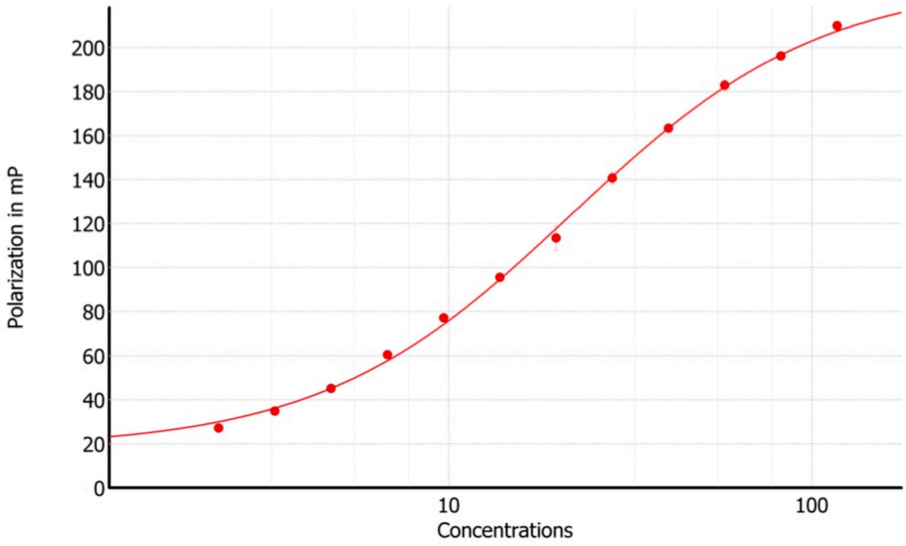


**4-Parameter fit**

Formula: y = Bottom + (Top - Bottom)/(1+(IP/x)^Slope)

| Parameter | Value |
|---|---|
| Top | 230.0 |
| Slope | 1.26032 |
| IC50 | 21.732 |
| log(IC50) | 1.337 |
| Bottom | 18.0 |
| r | 0.99944 |
| $r^2$ | 0.99888 |
| Curve Color: | |

**Figure S5.** Direct binding of increasing concentrations of EclA-C to fluorescently labelled fucoside **11** gave a $K_D$ of 21.7 $\mu$M. The titration was performed in 3 technical replicates.

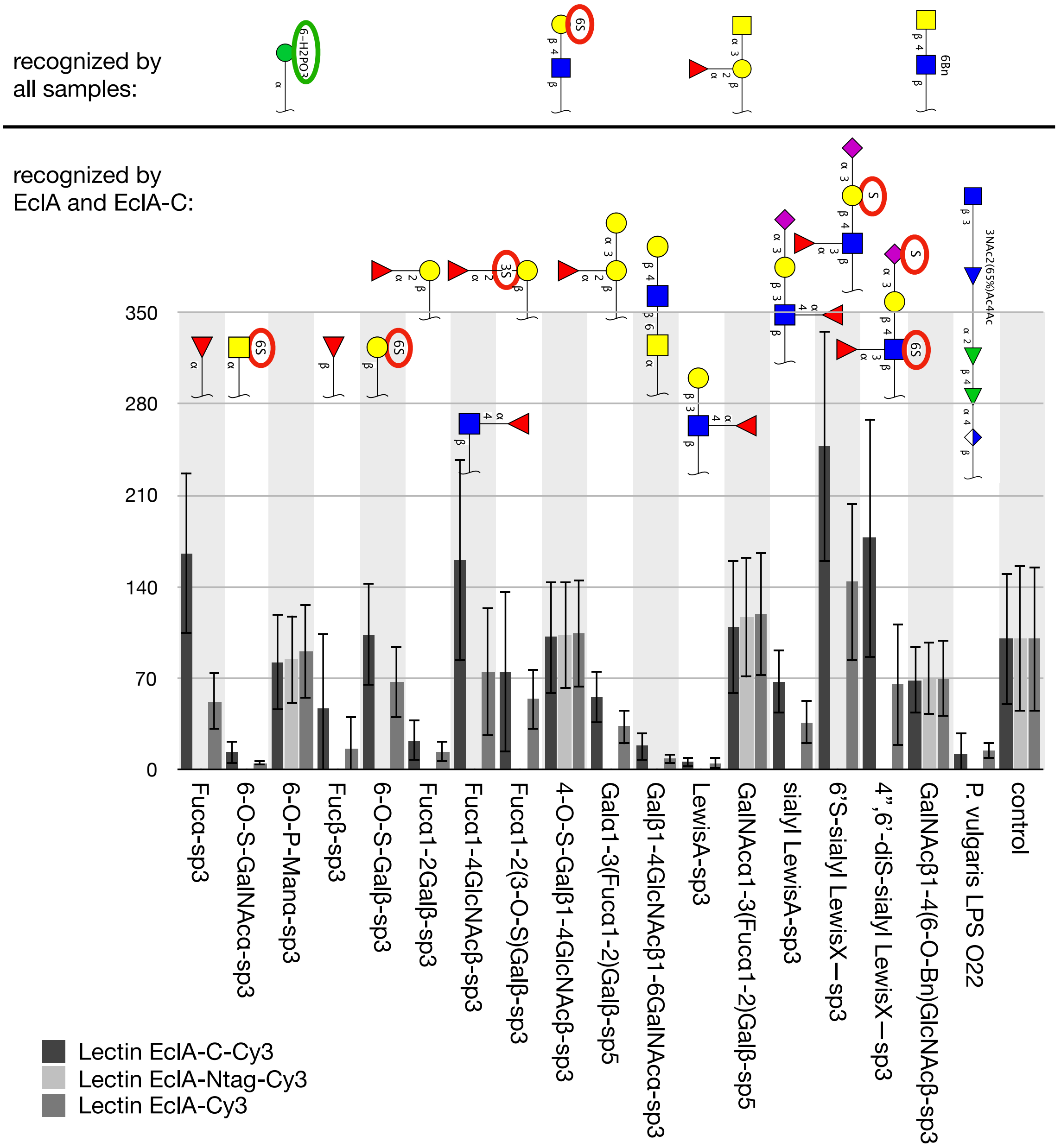


**Figure S6:** Analysis of Cy3-labelled EclA, EclA-N-tag and EclA-C on the Semiotik glycan array and comparative analysis of binding. The data confirmed fucose-, LewisA- and H-antigen-binding specificity for the C-terminus. In addition, a number of charged carbohydrates were bound by the constructs and four samples of which three lack fucose were equally recognized by all proteins, which contradicts CFG mammalian array and FP results. Depicted are all ligands above a fluorescence intensity threshold of 1000. y-axis is fluorescence intensity normalized to control.

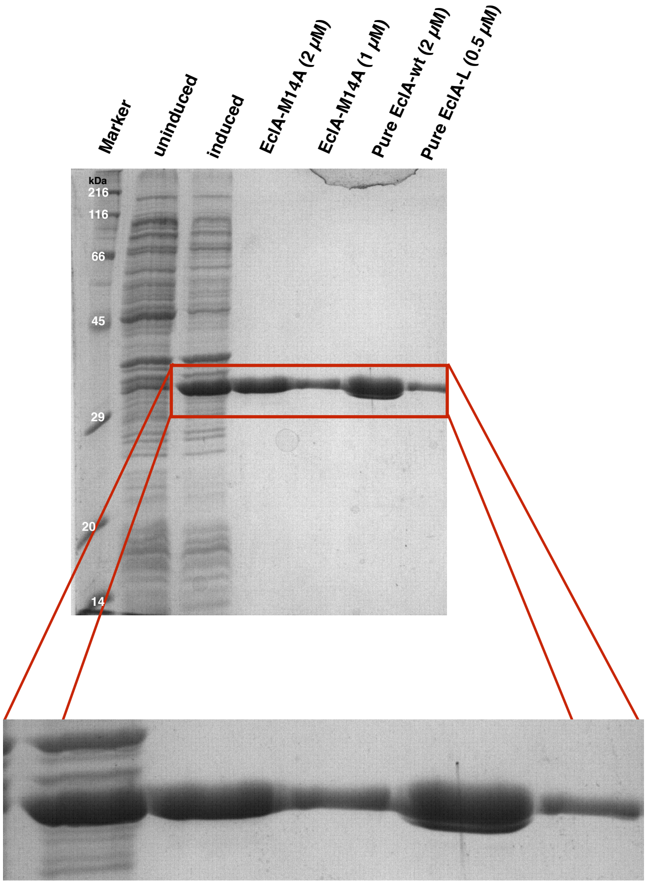


**Figure S7**: Expression and purification of EclA-L (EclA-M14A) shows homogenous protein species, compared to two close bands in EclA.

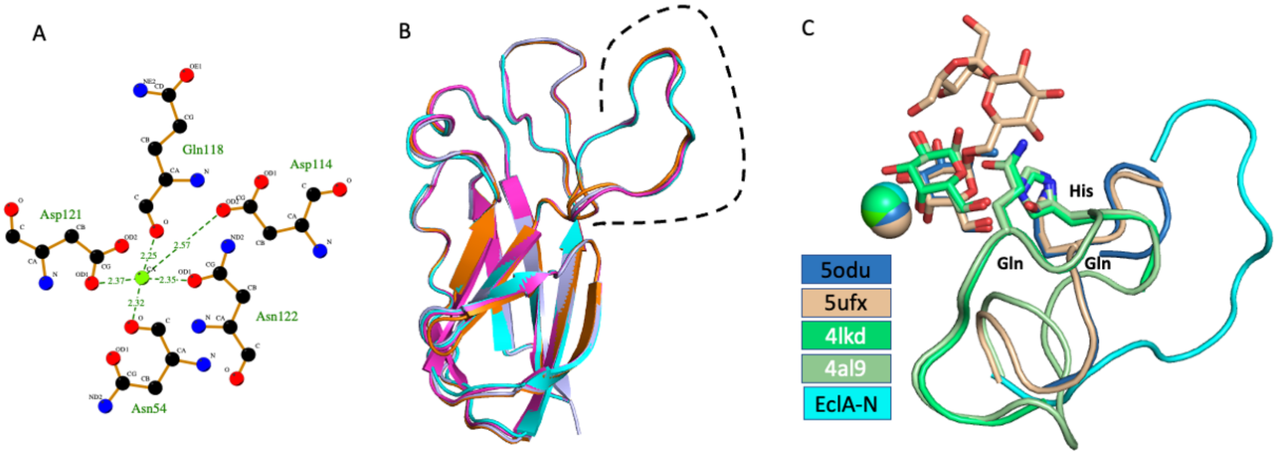


**Figure S8**: A. Calcium(II) coordination in EclA-N; B. Superposition of all four protomers of EclA-N in the asymmetric unit with the extended loop highlighted; C. Superposition of the binding site of EclA-N and its extended loop with galactose-containing structures of LecA (4al9, 4lkd) and PllA (5odu, 5ufx).

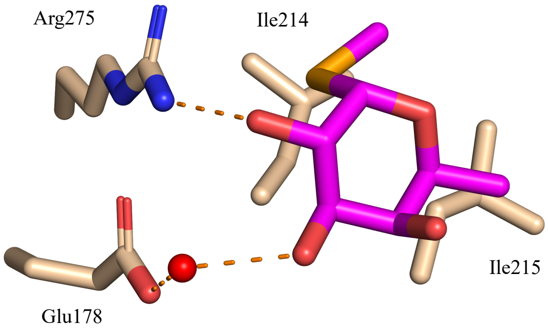


**Figure S9:** Schematic 2D representation of methyl α-L-selenofucoside interactions with residues of the symmetry mate. Hydrogen bonds are displayed in dotted orange lines, while all other residues exhibit hydrophobic interactions with the ligand. The water molecule is shown as a red sphere. For clarity only the interacting residues of the symmetry mate are shown as sticks.

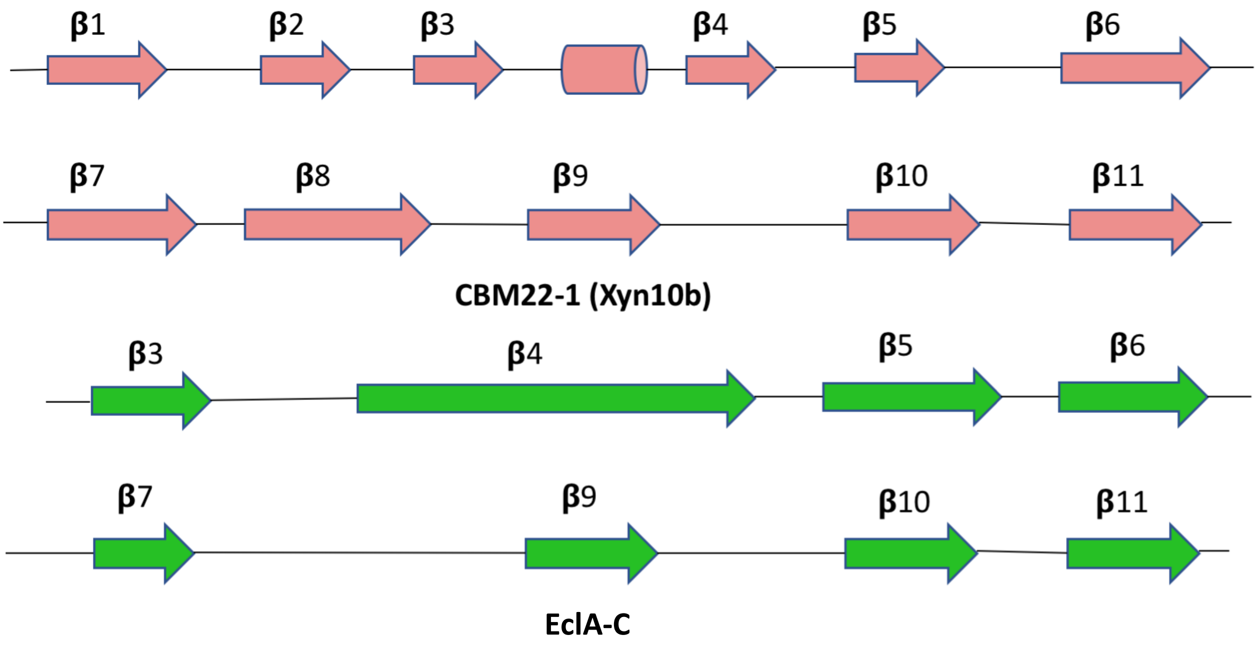


**Figure S10**: beta-strands in CBM22 compared to EclA-C.

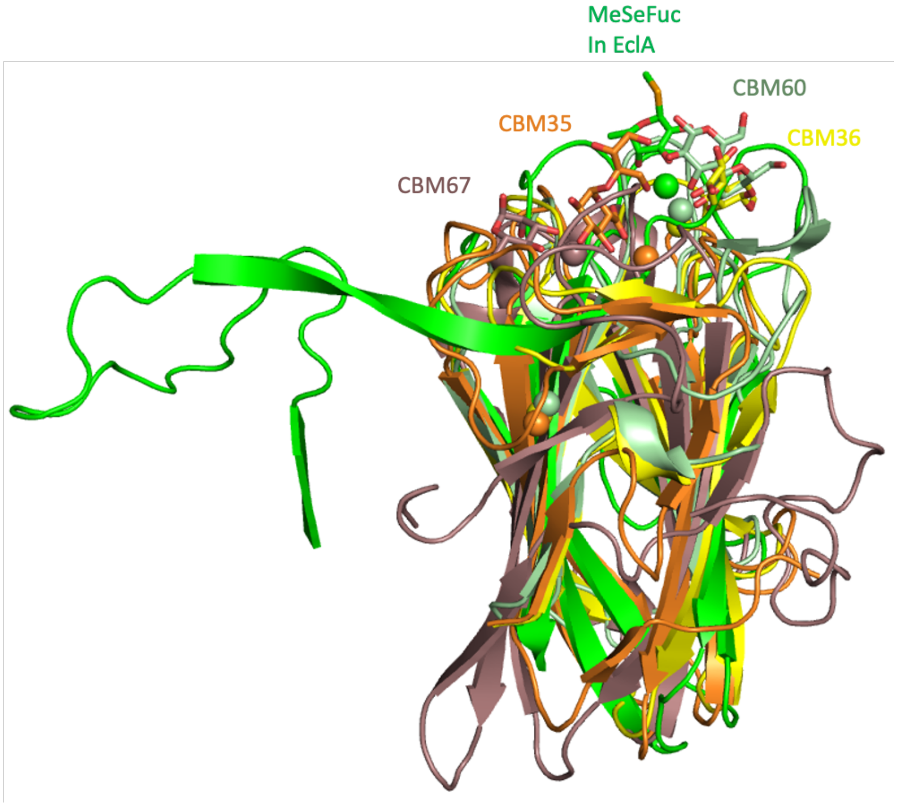


**Figure S11:** Superposition of the structures of EclA-C with CBM35, 36, 60 and 67 indicates calcium and ligand binding is not conserved.

**Table S1.** DNA Oligonucleotides used to amplify and sequence EclA-N-tag, EclA-C, EclA-L constructs by PCR.

| Primer | Sequence (5'- —> -3') | Restriction enzymes | His-tag |
|---|---|---|---|
| Forward EclA-N-tag (GB36) | GGAATTCCATATGCACCATCACCATCACCATGC GAAATTCTTCGCCACGCT | NdeI | ✓ |
| Reverse EclA-N-tag (GB37) | CGGGATCCTTACGCCGCGGCGGAAATA | BamHI | |
| Forward EclA-C (GB39) | GGAATTCCATATGGCAGCTCCGTTTGAAGATTT AAC | NdeI | |
| Reverse EclA-C (GB41) | CGGGATCCTTAACCCTTTATCTTCAGTTCCTTG | BamHI | |
| Forward EclA-L (M14A) (GB32) | CAGACGCGAGCGAAAATTTAATCTGGTC | | |
| Reverse EclA-L (M14A) (GB33) | TTCGCTCGCGTCTGACTCCTGTGAC | | |
| T7 promotor | TAATACGACTCACTATATAGG | | |
| T7 terminator | GCTAGTTATTGCTCAGCGG | | |

**Table S2**. Protein crystallography data collection and refinement statistics.

| | EclA-N-tag | EclA-C |
|---|---|---|
| **PDB ID** | 6YGQ | 6YF6 |
| **Data collection** | | |
| Space group | P 1 $2_1$ 1 | P $2_1$ $2_1$ $2_1$ |
| Cell dimension | | |
| a, b, c (Å) | 65.79, 50.46, 73.86 | 38.51, 63.67, 134.01 |
| a, b, g (°) | 90.00, 109.36, 90.00 | 90.00, 90.00, 90.00 |
| Wavelength (Å) | 1.00 | 0.967 |
| Resolution | 48.87 – 1.90 (1.94 – 1.90)* | 46.16 – 2.00 (2.05 – 2.00)* |
| Rsym or Rmerge | 0.066 (0.330) | 0.064 (0.321) |
| Rpim | 0.064 (0.300) | 0.018 (0.108) |
| CC (1/2) | 0.99 (0.88) | 1.00 (0.99) |
| I / sI | 8.0 (2.2) | 35.6 (8.5) |
| Completeness (%) | 99.9 (99.7) | 99.6 (96.3) |
| Redundancy | 3.4 (3.4) | 25.6 (18.4) |
| **Refinement** | | |
| Resolution (Å) | 40.20 – 1.90 | 36.57 – 2.00 |
| No. reflection | 36245 (3571) | 22932 (2175) |
| Rwork / Rfree | 0.182 / 0.225 | 0.193 / 0.241 |
| No. atoms | 4082 | 2471 |
| Protein | 3700 | 2200 |
| Ligands | 4 | 59 |
| Solvent | 378 | 240 |
| Protein residues | 480 | 280 |
| B-factors | 26.91 | 39.17 |
| Protein | 26.03 | 39.63 |
| Ligands | 35.54 | 29.08 |
| Water | 35.54 | 36.26 |
| R. m. s deviations | | |
| Bond length (Å) | 0.003 | 0.003 |
| Bond angels (°) | 0.63 | 0.60 |
| MolProbity score | 1.07 | 0.83 |
| *Values in parentheses are for highest-resolution shell | | |
